# From Toxicogenomics Data to Cumulative Assessment Groups: A Mechanistic Framework for Chemical Grouping

**DOI:** 10.1101/2025.01.24.634648

**Authors:** Sebastian Canzler, Julienne Lehmann, Jana Schor, Wibke Busch, Jörg Hackermüller

## Abstract

The grouping of chemicals based on shared properties or molecular mechanisms of action is pivotal for advancing regulatory toxicology, reducing data gaps, and enabling cumulative risk assessments. This study introduces a novel framework usingChemical-Gene-Phenotype-Disease (CGPD) tetramers derived from the Comparative Toxicogenomics Database (CTDbase). Our approach integrates toxicogenomics data to identify and cluster chemicals with similar molecular and phenotypic effects across diverse categories, including pesticides, pharmaceuticals, and industrial chemicals such as bisphenols and per- and poly-fluoroalkyl substances (PFAS). We validated our method by comparing CGPD Tetramer-based clusters with cumulative assessment groups (CAGs) for pesticides, demonstrating strong overlap with established groupings while identifying additional compounds relevant for risk assessment. Key examples include clusters associated with endocrine disruption and metabolic disorders.By bridging omics-derived molecular data with phenotypic and disease endpoints, this framework provides a comprehensive tool for chemical grouping and supports evidence-based regulatory decision-making, facilitating the transition to next-generation risk assessment methodologies.

## 1 Introduction

The growing complexity and volume of chemicals on the market, their tedious and year-long regulation procedures, and the increasing emphasis on reducing animal testing have spurred regulatory agencies and the scientific community to adopt alternative strategies, so-called New Approach Methods (NAMs) for chemical risk assessment^1^ (Westmoreland et al., 2022; Stucki et al., 2022; Schmeisser et al., 2023; European Commission and Cronin, 2024). Grouping chemicals based on common properties or molecular mechanisms of action has emerged as a powerful strategy for advancing safety evaluations, reducing data gaps, and generating scientifically informed insights for chemical risk assessment compliant with frameworks such as REACH^2^ (EU legislation on the Registration, Evaluation, Authorisation and Restriction of Chemicals). When applied correctly, grouping can minimize the need for experimental testing by leveraging data from structurally or functionally similar compounds (ECHA, 2023) complying with the 3R principles and the promotion of NAMs while facilitating read-across (RAx) methods or mixture risk assessment by accounting for additive or synergistic effects of chemicals using cumulative assessment groups.

The rapid development of omics technologies, such as transcriptomics, proteomics, and metabolomics, offers transformative potential for chemical grouping based on chemical-induced molecular responses of organisms or cells (Viant et al., 2024). Unlike traditional toxicology, omics assays capture the global biological response of a system to chemical exposure in a single experiment, providing insights into molecular mechanisms of action (Sauer et al., 2017; Canzler et al., 2020; Mortimer et al., 2022). Publicly curated resources such as the Comparative Toxicogenomics Database (`CTD`)^3^ have evolved to contain extensive omics-derived datasets (Davis et al., 2024).Using such resources and data allows the investigation of the molecular basis of toxicity and the associations of biomolecules and a wide range of chemicals with phenotypes and diseases.

The application of omics data to regulatory risk assessment has gained traction among agencies, e.g., the European Food Safety Authority (EFSA) and the European Chemicals Agency (ECHA), but also national institutions such as the US EPA or BfR in Germany, which recognize its benefits for chemical grouping, read-across, and cumulative risk assessment strategies (Marx-Stoelting et al., 2015; Harrill et al., 2019; EFSA et al., 2022b). EFSA, for instance, emphasized the role of omics technologies in transitioning to next-generation food and feed safety assessments, as outlined in their recently published roadmap on omics and bioinformatics (Radio et al., 2024), aligning with the EU’s Farm to Fork Strategy^4^ and Chemicals Strategy for Sustainability^5^ (CSS). Similarly, ECHA identified the use of omics methods as key elements in their roadmap for advancing NAMs to accelerate decision-making and reducing animal testing (ECHA, 2023).

In this context, grouping chemicals with diverse properties - such as pesticides, pharmaceuticals, or industrial chemicals such as bisphenols, or per- and poly-fluoroalkyl substances (PFAS), known for their widespread use and health impacts - has become a focus for environmental, consumer, and food safety assessment. Pesticides are widely used in agriculture but carry risks for human health and the environment, including neurotoxicity and endocrine disruption. While pharmaceuticals are essential for treating diseases, their environmental persistence and biological activity may pose unintended risks. Similarly, industrial chemicals, being ubiquitously present in consumer products, cosmetics, and food packaging, have been associated with adverse effects ranging from endocrine disruption to developmental and metabolic disorders (Lucas et al., 2022; Dalamaga et al., 2024; Chitakwa et al., 2024).

An important aspect of chemical safety evaluation is assessing the cumulative and synergistic effects of mixtures, particularly for chemicals with common toxicological effects and modes of action. EFSA and the European Council have recognized the need for methodologies that account for these combined effects, originally for pesticides, which are often encountered as mixtures in agricultural and environmental settings. In 2008, EFSA initiated efforts to develop cumulative risk assessment frameworks to summarize active substances into so-called Common/Cumulative Assessment Groups (CAGs) of pesticides (*EFSA*, 2008) which were related to the Common Mechanism Groups (CMG) that were established by the US EPA a few years prior (EPA, 2002). In 2012, an external scientific report to EFSA from the Danish Technical University identified toxicological effects and endpoints for grouping active pesticide substances into CAGs and proposed specific CAGs for consideration in Maximum Residue Limit (MRL) settings (Nielsen et al., 2012). Experimental evidence underscores the necessity of these approaches, demonstrating that mixtures of chemicals with common specific modes of action can interact additively or synergistically, producing effects greater than the sum of their individual impacts (Cedergreen, 2014). This finding reinforces the importance of identifying and grouping chemicals into CAGs, as it allows regulators to assess their cumulative risks more accurately and align safety evaluations with real-world exposure scenarios (Nielsen et al., 2012; Kortenkamp, 2022).

Despite the progress made, current methodologies face challenges in incorporating molecular-level data to improve the identification of CAGs and their combined effects. Omics technologies, which capture comprehensive molecular responses, offer a solution by enabling data-driven chemical grouping based on shared molecular effects. By linking omics data to phenotypic and disease endpoints, these approaches complement traditional risk assessments, providing deeper insights into mixture toxicology and cumulative risk.

Here, we present a novel methodology for grouping chemicals using CGPD tetramers (Chemical-Gene-Phenotype-Disease), which integrates curated chemical-based interactions from CTDbase (Davis et al., 2020). Unlike existing approaches focusing on structural or functional similarities, our framework leverages multi-dimensional toxicogenomics data to identify shared molecular mechanisms and phenotypic effects across diverse chemical classes, facilitating their joint consideration in cumulative risk assessments or read-across safety evaluations.

Our study addresses modern regulatory needs by expanding the scope of cumulative risk assessments to include previously unrecognized chemicals, offering a robust, data-driven tool to enhance safety evaluations. By bridging molecular mechanisms and regulatory frameworks, it supports next-generation risk assessment methodologies, advancing regulatory toxicology and informing evidence-based decisions while addressing challenges in data quality, filtering, and methodological refinement.

## 2 Material and Methods

### 2.1 CTD Nomenclature

In the context of this study, we adopt the term interaction as it is used in the CTDbase to describe relationships between chemicals and biological entities, such as genes or phenotypes. It is important to note that interaction in `CTD` does not necessarily imply a direct physical binding or biochemical interaction, but rather an association curated from experimental studies and literature. We also use the term association interchangeably with interaction to indicate links between chemicals and their reported biological effects. Therefore, our terminology aligns with `CTD`’s annotation format, specifically referring to curated data tables such as chemical-gene interactions, chemical-phenotype interactions, or chemical-disease associations which serve as the basis for our methodology.

### 2.2 Database Generation

Processed omics data from the Comparative Toxi-cogenomics Database (`CTDbase`) and chemical properties from `PubChem` were compiled into a SQLite database with minor modifications. Interaction tables from `CTDbase` were downloaded in TSV format, columnheaders were standardized to lowercase and stripped of ‘MESH:’ prefixes in MeSH ID columns for consistency. The processed files were then converted into database tables named after their source files.

`CTD` relies on MeSH IDs to identify chemicals and diseases while `PubChem` assigns chemicals a unique compound identifier (CID). The Medical Subject Headings (MeSH) is a controlled and hierarchically organized vocabulary produced by the National Library of Medicine. To map MeSH IDs to `PubChem` CIDs, we utilized the `PubChem` Identifier Exchange Service^6^. Of 14 489 MeSH IDs from `CTD`-annotated chemicals with curated chemical-gene interactions, 12 307 (85%) were successfully mapped to CIDs.

For each chemical with a CID we extracted data from `PubChem`, such as InChI key, SMILE, molecular mass, chemical formula, and external identifiers. Two database tables were created: one for chemical descriptions and one for chemical names and external IDs.

The SQLite database finally comprises 18 tables — 16 from `CTDbase` and two from `PubChem`. Download date was March 20, 2023. Details of the tables and data sources are provided in Supplementary Tables S1 and S2. The database dump was uploaded to Zenodo^7^.

#### Gene mapping

`CTDbase` uses primary gene IDs as gene identifiers in chemical-gene interactions, which may not always align with organism-specific Entrez Gene IDs. To address this, a species-specific gene ID column was added to the database. Mapping was performed using NCBI’s `gene_orthologs`^8^ file, which provides pairwise orthologs. For each chemical-gene interaction, we verified if the primary gene ID matched the annotated organism for which the interaction was curated. If not, the correct species-specific ID was identified via the `gene_orthologs` file and updated in the database. Interactions lacking orthologous genes for the target species were excluded.

### 2.3 Workflow for Chemical Grouping

#### 2.3.1 Calculation and Filtering of CGPD Tetramers

A CGPD tetramer is a computationally predicted information block describing a chemical (C) that interacts with a gene product (G), inducing a non-disease phenotype (P) linked to a disease (D), as shown in Figure 1(A). It integrates five evidence statements: four from `CTDbase` and one inferred via GO enrichment analysis from NCBI.

**Fig. 1:**
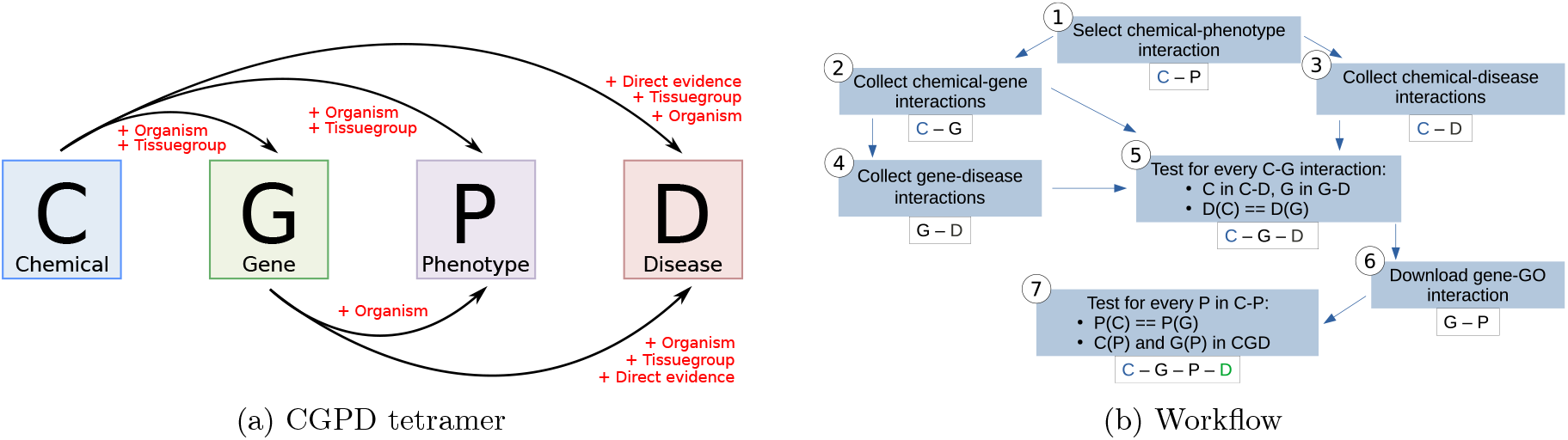
Building of a CGPD tetramer by integrating five different interactions. (A) CGPD tetramer strutcure and filter options. Chemical - gene interactions (C-G), chemical - phenotype associations (C-P), chemical - disease associations (C-D), and gene - disease associations (G-D) are taken from `CTDbase`. Gene - phenotype associations (G-P) were collected from NCBI. Filter options for individual associations are indidcated in red. (B) Summarized workflow to derive CGPD tetramers based on processed omics data from `CTDbase`.

The basic concept was adapted form Davis et al. (2020) and Grondin et al. (2021) with modifications for optional filtering steps. The workflow is depicted in Figure 1(B) and outlined as follows:

First, the initial set of chemicals (*C*) was compiled from annotated chemical-phenotype interactions in `CTDbase` (Step 1). For each chemical *c* ∈ *C*, associated chemical-gene and chemical-disease interactions were extracted (Step 2 & 3). Genes identified in chemical-gene interactions were used to gather gene-disease associations (Step 4).

Second, CGD Triplets were created by identifying shared associations between a chemical, a gene, and a disease (Step 5).

Next, the ncbi2go annotation file^9^ was utilized to collect gene-phenotype interactions for all genes present in the CGD Triplets (Step 6). Interactions were identified by leveraging the species-specific Entrez gene IDs from the described mapping procedure.

Finally, the CGPD tetramers were assembled (Step 7) by integrating the phenotype-related information from both chemical-phenotype and gene-phenotype interactions with the CGD Triplets. Phenotypes were included if they were linked to both the chemical and the gene in the CGD Triplet.

#### 2.3.2 Filtering Options

To enhance the precision of CGPD tetramer calculations, we implemented several filtering options that can be applied individually or in combination. These filters are adaptations of the overall workflow described above, allowing for more stringent data selection. The specific filters and their application within the CGPD tetramer calculation are illustrated in red in Figure 1(A).

##### Filter for organisms

Chemical-gene and chemicalphenotype interactions in `CTDbase` are explicitly annotated with organism information, allowing species-specific filtering in Steps 1 and 2. Gene-phenotype interactions are already linked to phenotypes via species-specific gene IDs, hence no further filtering is required. Disease-related interactions, however, lack the annotation with organism information. To address this, PubMed IDs from chemical-disease and gene-disease interactions were crossreferenced with chemical-gene interactions to determine organism-specific information, enabling linkage of disease interactions to the organisms of interest.

##### Filter for tissue groups

Chemical-phenotype interactions are annotated with anatomy terms, which form a subset of descriptors from the ‘Anatomy’ category in the MeSH vocabulary. Individual anatomy terms were grouped into so-called tissue groups to reflect more coarse-grained tissues and structures enabling tissue-specific filtering. In total, 28 tissue groups were defined. 25 tissue groups were originally mentioned by Nielsen et al. (2012), while three additional groups were formed: Breast, Fat tissue, and Pluripotent stem cells. Supplementary Section S0.1 lists the groups with their 506 assigned anatomy terms.

Amongst all `CTD`-listed anatomy terms, 38 are related to cancer, tumor, or carcinoma. Those terms may not correspond to a particular tissue group, e.g., HeLa cells, while other terms commonly describe organ-specific models, e.g., Hep G2 cells as human hepatocyte model. To account for this discrepancy, the possibility to exclude all cancer/tumor/carcinoma related anatomy terms from CGPD tetramer inference was implemented.

Chemical-gene interactions lack anatomy terms but include Pubmed IDs. These IDs, collected from chemical-phenotype interactions in Step 1, enable indirect tissue group filtering in Step 2.

Chemical-disease and gene-disease interactions are not linked to anatomy terms, even though they may be highly tissue-specific. Therefore, a mapping procedure was implemented linking diseases to anatomy terms and hence tissue groups. `CTD`-annotated disease names, their synonyms, and parent disease names were used to manually match existing anatomy terms. Diseases may be linked to one or multiple tissue groups based on such a match; otherwise, they default to all tissue groups. Broader disease terms were mapped to overarching categories linked to multiple tissue groups, such as:

– **Endocrine system** Adrenal gland, Pituitary gland, Thyroid gland, Parathyroid gland
– **Urogenital diseases** Urinary bladder, Kidney, Reproductive system
– **Urologic diseases** Urinary bladder, Kidney

The mapping file containing the disease - tissue group associations is uploaded to Zenodo^10^.

##### Direct evidence for associations with diseases

Diseaserelated associations in `CTD` (C-D or G-D) are categorized as ‘direct evidence’ when they are manually curated from published literature. They are further classified as ‘marker/mechanism’ or ‘therapeutic’.

‘Marker/Mechanism’ indicates a chemical or gene is associated with the disease’ etiology, such as chemical exposure causing lung cancer or a gene mutation causing liver cancer. ‘Therapeutic’ refers to chemicals with known/potential therapeutic roles or genes that are therapeutic targets in the disease treatment.

‘Inferred’ disease associations are not manually curated from published literature but rather based on shared entities: either through a common gene (chemical-disease interaction) or a common chemical (gene-disease interaction).

The evidence type (‘marker/mechanism’, ‘therapeutic’, or ‘inferred’) is listed in the respective database table and can be used for filtering. By default, CGPD tetramers include only disease interactions with direct evidence, though users may filter by specific evidence types or disable this filtering option.

#### 2.3.3 Grouping of Chemicals

To group chemicals according to common biological effects, two grouping strategies were applied using the previously established CGPD tetramers: (1) GPD-based grouping and (2) PD-based grouping.

##### GPD-based grouping

Chemicals were grouped if they shared common GPD trimers capturing gene, phenotype, and disease information. For example, Carbendazim and Bisphenol A were grouped because they interact with the gene *Esr1*, associated with the phenotype steroid metabolic process, and the disease Oligospermia.

##### PD-based grouping

Chemicals were grouped if they share common PD dimers capturing phenotype and disease information but excluding gene information. For example, Carbendazim and Fenvalerate were grouped despite interacting with different genes (*Esr1* and *Star*, respectively), as they share the common phenotype steroid metabolic process and the disease Oligospermia.

In both strategies, the grouping was conducted separately for tissue groups and target organisms.

#### 2.3.4 Clustering of Chemical Groups

Two approaches were used to cluster chemical groups:

(1) clustering by the semantic similarity of associated phenotypes and (2) clustering by the similarity of associated list of chemicals.

##### Semantic similarity of phenotypes

The semantic similarity of phenotypes, based on Gene Ontology (GO) annotations, was computed using the pygosemsim python package. All CGPD-annotated phenotypes were collected and their pairwise semantic similarity was calculated using the graph-based ‘Lin’ measure. This measure calculates similarity by considering the information content (IC) of the most informative common ancestor of the compared phenotype terms (Pesquita et al., 2009).

Using the resulting similarity matrix, connected components of phenotypes with a pairwise similarity above a particular threshold were identified. In the GPD-grouping strategy, similar phenotypes were merged if they shared common gene and disease IDs. In the PD-grouping, similar phenotypes were merged if they shared a common disease ID.

##### Clustering on chemical lists

The similarity of chemical groups was evaluated by calculating the Tanimoto similarity of their associated lists of chemical. The Tanimoto similarity between two groups *C*_*i*_ and *C*_*j*_ is defined as:

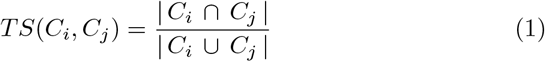

where *C*_*i*_ and *C*_*j*_ are the chemical lists associated with group *i* and *j*, respectively.

A Tanimoto similarity matrix was generated and connected components of chemical groups with pairwise similarity above a user-defined threshold were identified.

#### 2.3.5 Applied Filtering and Grouping Strategies

Separate filtering for three target organisms (human, mouse, and rat) and 28 tissue groups (see Supplementary Section S0.1) was applied for chemical-gene (C-G), chemical-phenotype (C-P), and gene-phenotype (G-P) interactions. Additionally, for the evaluation of the grouping and clustering procedure the following five different combinations of filters for disease-related interactions were implemented:

(v1) Direct evidence for ‘marker/mechanism’, tissuespecific diseases, species-specific diseases

(v2) Direct evidence for ‘marker/mechanism’ and ‘therapeutic’, tissue-specific diseases, speciesspecific diseases

(v3) Direct evidence for ‘marker/mechanism’ and ‘therapeutic’, tissue-specific diseases

(v4) Direct evidence for ‘marker/mechanism’ and ‘therapeutic’, species-specific diseases

(v5) Direct evidence for ‘marker/mechanism’ and ‘therapeutic’

CGPD tetramers generated under these filtering strategies were grouped in two ways: based on common genes, phenotypes, and diseases (GDP-grouping), and based on common phenotypes and diseases (PD-grouping). These groupings were further clustered using semantic similarity of phenotypes and/or Tanimoto similarity of their chemical lists at three thresholds (0.5, 0.75, 0.9). In total, 150 groupings and clusterings were generated for each of the three organisms, encompassing all combinations of filtering, grouping, and clustering methods.

### 2.4 Implementation and Data Availability

The complete workflow has been implemented in Python, providing a robust and flexible platform for chemical grouping and clustering analyses. It was designed with two key functionalities accessible through subcommands: The **download** subcommand automates the retrieval of required datasets from `CTDbase` and `PubChem`, subsequent processing, formatting, and database construction. The **grouping** subcommand facilitates the generation and analysis of CGPD tetramers, including the application of various filtering, grouping, and clustering methodologies. Parameters can be customized in a YAML configuration file, allowing for seamless adaptation to different contexts.

To promote accessibility and reproducibility, the workflow is packaged as a Docker container hosted at the Helmholtz Codebase. The underlying Python code is open-source, distributed under the CC-BY License, and is available in the project’s GitHub repository^11^.

### 2.5 Chemical Classes

#### Pesticides and Cumulative Assessment Groups

Documents from the European Food Safety Authority (EFSA) were utilized to compile lists of pesticides and their classification into Cumulative Assessment Groups (CAGs). The foundational work by Nielsen et al. (2012), commissioned by EFSA, proposed a tiered approach to identify CAGs for cumulative risk assessment (CRA), ranging from broad groups affecting specific organs or tissues (Level 1) to chemicals with a common mechanism of action (Level 4). This report categorized 248 pesticides, of which 211 were successfully mapped to MeSH IDs. EFSA has since developed specific CAGs for the nervous system (EFSA et al., 2019b), thyroid gland (EFSA et al., 2019a), and craniofacial alterations (EFSA et al., 2022a). These groups included 125, 133, and 51 pesticides that were mapped to MeSH IDs, respectively.

The EFSA PARAM catalogue version 7^12^ was also used to identify pesticides. Filtering this data set yielded 1755 compounds, of which 856 were mapped to MeSH IDs.

Additionally, 327 compounds were collected from `CTDbase` by identifying pesticide-related terms in their MeSH entries. A detailed list of these terms and the corresponding number of pesticides is provided in Supplementary Table S3.

In total, 938 unique pesticides were consolidated from these sources, and an overlap analysis of the data sets is presented in Supplementary Figure S1.

#### Pharmaceuticals

Pharmaceutical data were obtained from two repositories: the DrugBank academic release version 5.1.11^13^ and `CTDbase`. The DrugBank dataset contained 16 575 drugs, of which 7590 compounds were retained after cross-referencing with `CTDbase` using drug names and CAS numbers.

From `CTDbase`, pharmaceuticals were identified using the MeSH dictionary to extract relevant pharmacological terms, resulting in 960 compounds. The list of MeSH terms is provided in Supplementary Table S4.

Compounds previously classified as pesticides were excluded from both datasets, resulting in 7447

DrugBank pharmaceuticals and 932 from `CTDbase`, for a total of 7854 unique pharmaceuticals.

#### Industrial chemicals

Classifying non-pharmaceutical and non-pesticide compounds into specific functional groups is challenging, but three key chemical groups relevant to regulatory applications were selected: phthalates, bisphenols, and per- and polyfluoroalkyl substances (PFAS).

Bisphenols were identified using `CTD`-annotated names and synonyms by searching for the ‘bisphenol’ pattern, resulting in 45 compounds. Similarly, phthalates were identified using the ‘phthalate’ pattern, yielding 113 compounds. PFAS, a more heterogeneous group, were identified through their parent MeSH ID (Fluorocarbons, MeSH ID: D005466), resulting in a set of 450 compounds.

## 3 Results and Discussion

### 3.1 Data Coverage in CTD

Before evaluating the proposed grouping method, we analyzed the available data in the `CTDbase` to assess its coverage across organisms, tissues, and chemical classes. This analysis identified key patterns and limitations, providing a foundation for tailoring, filtering and grouping strategies to the dataset’s structure.

#### 3.1.1 Organism- and Tissue-specificity

Data from `CTDbase` revealed a total of 625 organisms with nearly 1.7 million annotated chemical-gene interactions, of which 26 species had more than 1000 interactions. The highest numbers of chemical-gene interactions were observed for human (693 470), mouse (420 453), and rat (342 393), followed by zebrafish with fewer than 80 000 interactions. All remaining organisms combined for nearly 130 000 interactions, see Figure 2 A.

**Fig. 2:**
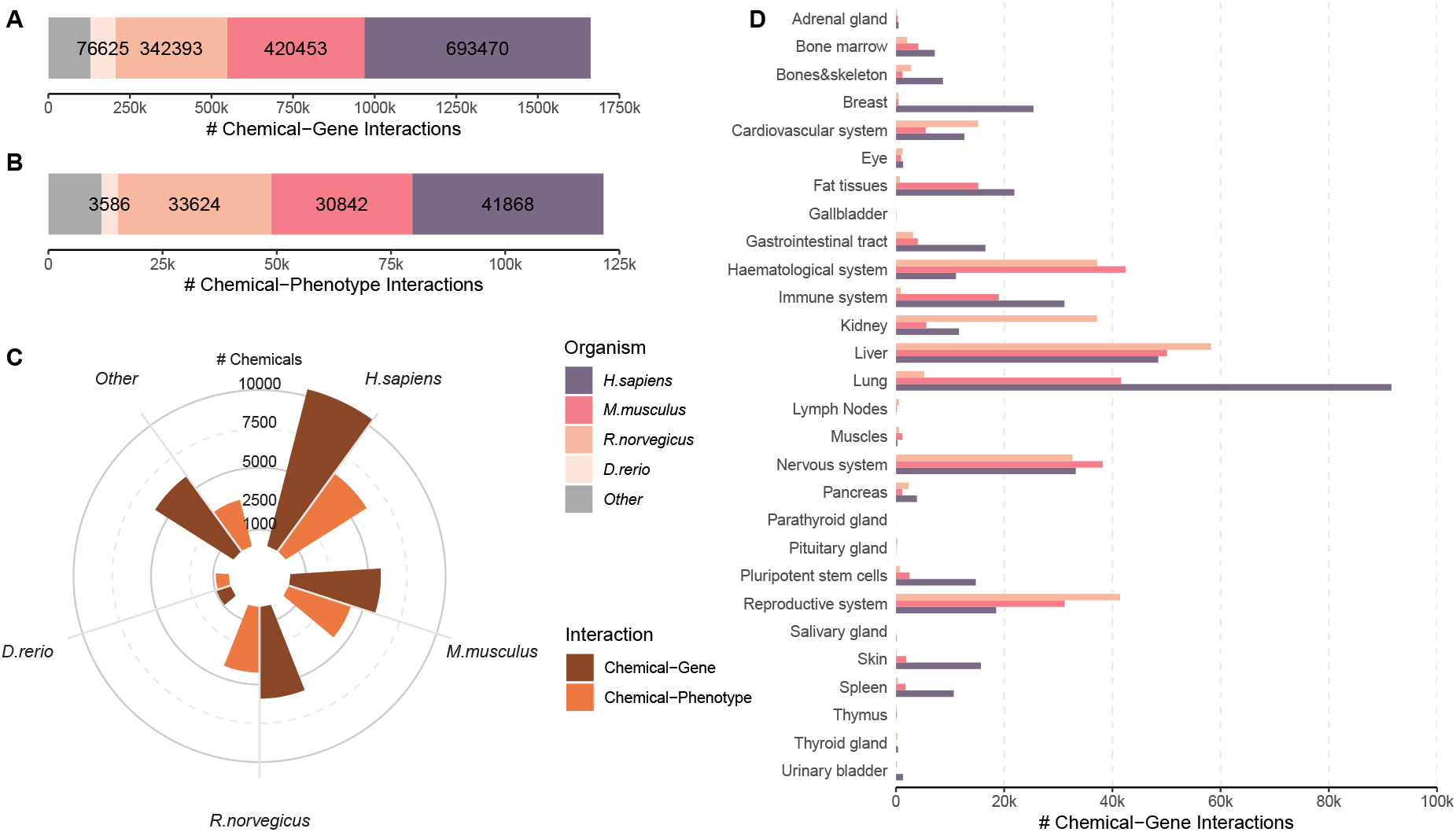
Overview of CTD data availability. (A) Number of annotated chemical-gene interactions for specific organisms. The top 4 organisms are shown as individual entities, all other organisms are combined. (B) Number of annotated chemical-phenotype interactions for specific organisms. Top 4 organisms are shown individually, other species are combined (see color legend). (C) The number of distinct chemicals in chemical-gene and chemical-phenotype interactions is shown for the top 4 organisms individually, all other species are combined. (D) Number of annotated chemical-gene interactions for the 28 tissue groups and the three organisms human, mouse, and rat.

Similar patterns were observed for chemicalphenotype interactions. Slightly over 121 000 C-P interactions were distributed across 346 species. Humans again had the highest number of interactions (41 868), followed by rat (33 624), mouse (30 842) and zebrafish (3586). The remaining organisms combined for a total of 10 000 chemical-phenotype interactions, see Figure 2 B.

For humans, 10 441 different chemicals were contained in chemical-gene interactions, while 6188 chemicals were listed in chemical-phenotype interactions. For rats and mice, we found comparable numbers, with 5909 and 5824 chemicals in chemical-gene interactions and 4226 and 4205 in chemical-phenotype interactions, respectively, see Figure 2 C.

Notably, approximately 60% of the chemicals were shared between rat and mouse in both interaction types. Fewer than 1000 different chemicals were annotated in zebrafish, while all remaining organisms accounted for 5879 and 3773 chemicals in chemical-gene and chemical-phenotype interactions, respectively.

The data clearly highlights discrepancies in the number of annotated interactions and chemicals across various species.

Rat (512), mouse (506), and human (476) were also the species with the highest number of anatomy terms in chemical-phenotype interactions. Human data were enriched with cell line-based anatomy terms (e.g., Hep G2, MCF-7), while rodent data predominantly involved whole organs (e.g., liver, lung, brain, heart) and fluids (e.g., serum, blood, urine).

An overview of chemical-gene interactions distributed across tissue groups is shown in Figure 2 D. Similar to the anatomy terms, we also found species-specific patterns. By far the most chemical gene interactions in human were curated for lung tissue followed by liver, and the nervous system. In rats, most interactions were collected for liver, the reproductive, and the hematological system. Mouse chemical-gene interactions were predominantly curated for liver, the repoductive system, and kidney. These observations underscore the variability in available curated data across different tissue groups and species. To capture specific molecular effects and associations with genes, phenotypes, or diseases, it is essential to filter for individual target tissues and organisms. Focusing on the three organisms - human, mouse, and rat - we implemented species-specific and tissue-specific filtering strategies to address these differences and enhance the specificity of the analysis.

#### 3.1.2 Coverage of Chemical Classes

To evaluate the representation of specific chemical classes within CGPD tetramers, we analyzed their coverage under different filtering strategies applied to disease-related interactions. This analysis focused on chemical groups with high regulatory relevance, including pesticides and pharmaceuticals, as well as industrial chemicals such as bisphenols, phthalates, and per- and polyfluoroalkyl substances (PFAS), which are known to be associated with adverse health effects, including endocrine disruption and developmental abnormalities (Alamri et al., 2021). Their presence in food contact materials and potential for migration into food have raised significant public health concerns (Pedersen et al., 2008).

A comparison of the most stringent filtering strategy (v1) with the most relaxed one (v5) is shown in Figure 3; a complete overview of all filtering strategies is provided in Supplementary Figure S2. In principle, the representation of chemical classes in CGPD tetramers varied substantially. Pesticides accounted for a smaller proportion of tetramers compared to other chemical classes, representing 7% of tetramers in rats under filtering strategy v1, decreasing to 5% in v5. Similar trends were observed in humans and mice, see Figure 3 A. The number of distinct pesticides reflected in these tetramers was comparable across the three species, ranging from 52–66 in v1 to 69–90 in v5, constituting 6–10% of all chemicals, cf. Figure 3 B.

**Fig. 3:**
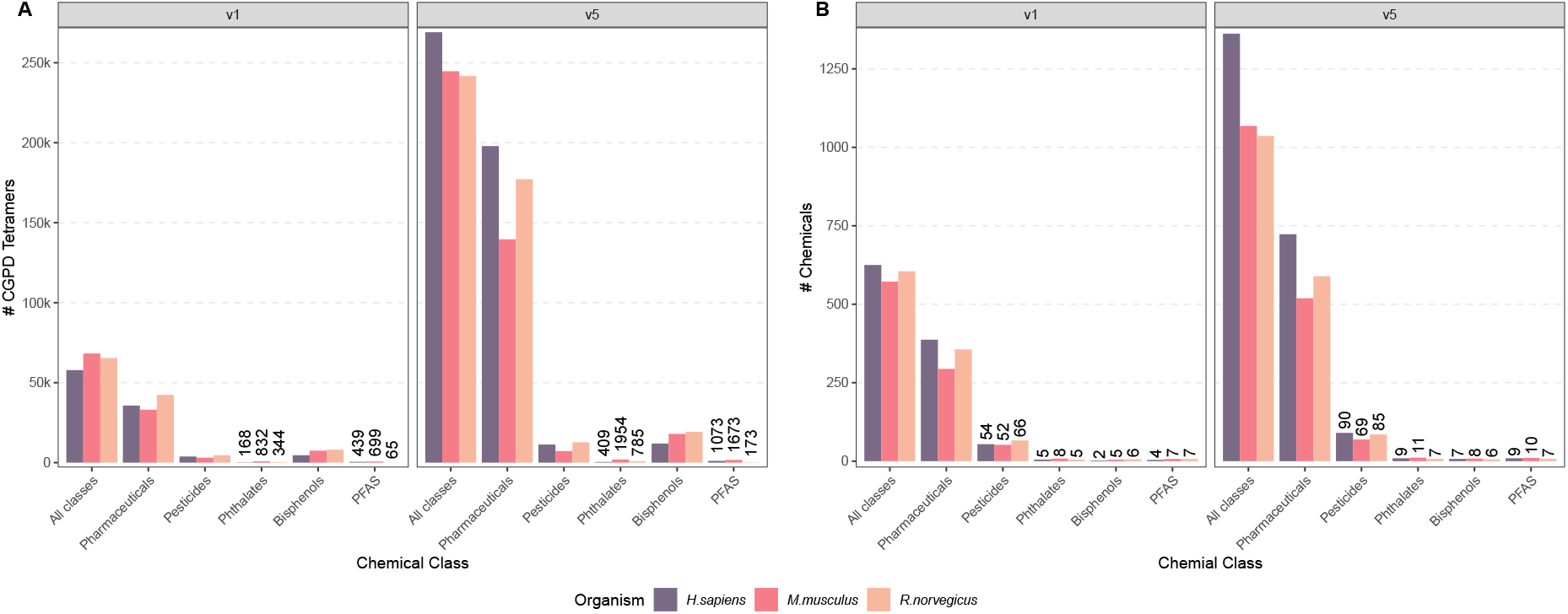
Subsetting CTD-derived data to different chemical classes. (A) Number of CGPD tetramers calculated from `CTDbase`. (B) Number of chemicals for which CGPD tetramers could be calculated. tetramers have been individually calculated for the three target organisms human, rat, and mouse. For visibility reasons we plotted here only the most stringent (v1) and most relaxed filtering strategies (v5). All five strategies can be seen in Supplementary Figure S2. Pesticides have been collected from the EFSA PARAM catalogue (denoted as EFSA Pesticides), from four different CAG reports (EFSA CAGs) and the `CTDbase` itself based on the MeSH dictionary (`CTD` Pesticides). Pharmaceuticals have been collected from the Drugbank and the `CTDbase` itself based on the MeSH dictionary. Bisphenols, Phthalates, and PFAS have been collected from `CTDbase` using the MeSH dictionary.

Pharmaceuticals were the most represented chemicals, reflecting the rich data coverage in `CTDbase`. In filtering strategy v5, pharmaceuticals accounted for 73% of tetramers in humans and rats and 57% in mice. The proportion of unique pharmaceuticals involved in tetramer calculations was also high: 53% in humans, 57% in rats, and 48% in mice.

Other chemical classes, such as bisphenols, phthalates, and PFAS, showed distinct patterns. For example, Bisphenols were prominently represented in terms of tetramers, with nearly 20 000 in rats under filtering strategy v5. However, the number of unique bisphenols was limited, ranging from 2-9 compounds depending on the filtering strategy and organism. Phthalates and PFAS, while generating fewer CGPD tetramers overall, had comparable representation in terms of unique compounds, with up to 11 individual chemicals in the mice data in filtering strategy v5.

A detailed distribution of tetramers and chemical classes across tissue groups is provided in Supplementary Figures S3 and S4. There, pesticides showed consistently the highest number of tetramers associated with the nervous system. High numbers of pesticide-related tetramers were found in the reproductive system and liver in the rat data set, while human data showed greater representation in the lung and gastrointestinal tract. In contrast, pharmaceutical data displayed broad tissue coverage, with the liver being a dominant tissue group in humans and mice, whereas the nervous system had the highest representation in the rat data set. Bisphenols were primarily associated with the reproductive system in rats, mice, and humans, with additional relevance in the liver, breast, and lung depending on the species. Details are provided in Supplementary Section S1.1.

These results demonstrate significant variability in CGPD tetramer coverage across chemical classes, tissue groups, and species, highlighting the importance of filtering strategies and organism-specific data for assessing molecular effects.

### 3.2 Comparison of Grouping/Clustering Strategies

To assess the effectiveness of different parameter combinations for grouping and clustering chemicals, we analyzed 30 configurations using the most stringent filtering strategy (v1). The analysis varied grouping criteria (GPD-based vs. PD-based) and clustering thresholds for semantic and Tanimoto similarity to explore their ability to identify biologically relevant molecular relationships. Results were compared across three target organisms, with a graphical summary presented in Figure 4.

**Fig. 4:**
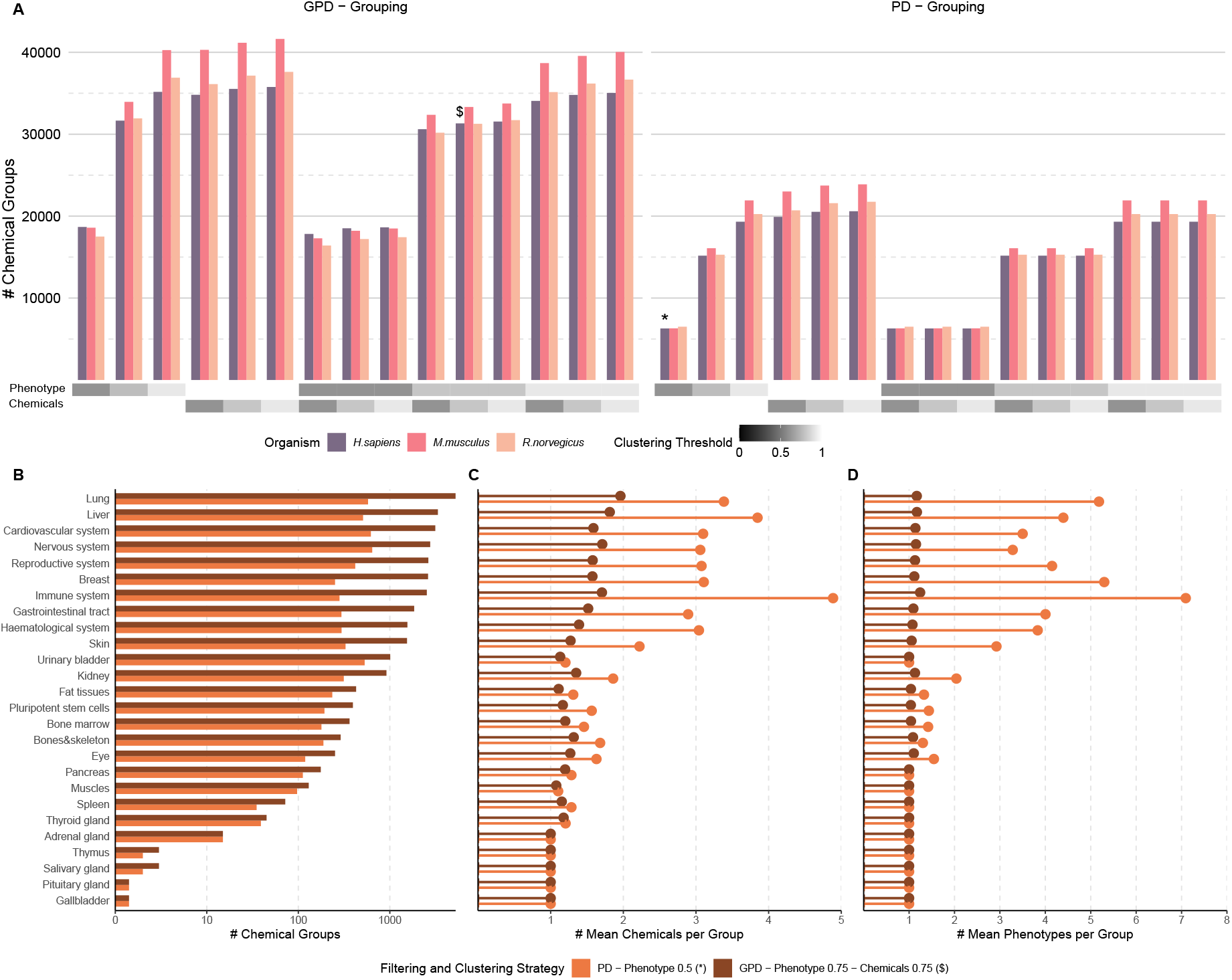
Visualizing differences in grouping and clustering strategies. (A) Number of chemical groups that have been formed from CGPD tetramers based on different grouping (PD vs GPD) and clustering strategies (phenotype and chemical-based clustering). The most stringent filtering version v1 was used to collect the CGPD tetramers. (B-D) Comparison of two particular grouping and clustering approaches with respect to different tissue groups in human. First, a very stringent method based on PD grouping and phenotype-based clustering with a threshold of 0.5 (marked with * in Subfigure A). Second, a grouping strategy utilizing GPD grouping and a clustering based on phenotypes and chemicals with more relaxed thresholds of 0.75 (marked with $ in Subfigure A). (B) Comparison of the number of chemical groups in different tissues. (C) Comparison of the mean number of assigned chemicals in individual tissue groups. (D) Comparison of the mean number of assigned phenotypes in different tissue groups.

PD-based grouping resulted in a drastically reduced number of chemical groups compared to GPD-based grouping, as shown in Figure 4 A. This reduction is attributed to the omission of commonly targeted genes in the grouping process, leading to groups with more chemicals included. The pattern was consistent across humans, mice, and rats.

Clustering strategies had varying effects. Clustering based on the Tanimoto similarity of assigned chemical lists had a marginal impact, even at a low threshold of 0.5. In contrast, clustering by phenotype similarity had a substantial effect on the number of chemical groups. For example, GPD-grouping with phenotype-based clustering at a threshold of 0.9 yielded approximately 35 000 chemical groups, which decreased to 18 000 groups at a threshold of 0.5 for human data. This indicates that phenotype-based clustering effectively consolidates chemical groups, resulting in fewer but larger clusters with broader chemical and molecular diversity. Similar trends were observed when both clustering strategies were combined, with phenotype clustering having the dominant impact.

A direct comparison of two human-based cases highlighted the impact of parameter choices on chemical grouping. GPD-grouping with clustering thresholds of 0.75 for both phenotypes and chemical lists resulted in a high number of groups (marked with $ in Figure 4 A), On the contrarty, PD-grouping with phenotype clustering at a threshold of 0.5 produced substantially fewer groups (marked with * in Figure 4 A). Comparing the distribution of chemical groups across tissue groups (Subfigure B), a clear difference was observed in information-rich tissues such as the liver, lung, and immune system. Consequently, these tissues showed also the largest differences in the mean number of assigned chemicals and phenotypes, shown in Figure 4 C and D, respectively.

In summary, the parameter choices for grouping and clustering significantly influence the number and size of chemical groups. PD-grouping produces fewer but larger groups of chemicals, while GPD-grouping generates more groups with fewer assigned chemicals. These differences are most pronounced in data-rich tissues heavily represented in `CTDbase`. Clustering by phenotype similarity exerts a greater effect than clustering by chemical list similarity and provides a mechanism to control group size and phenotypic diversity. Lower phenotype clustering thresholds lead to greater phenotypic variety within groups, though all phenotypes remain linked to the same disease through the CGPD tetramers. This flexibility enables tailored analyses based on the specific aims of the study.

### 3.3 Tissue-specific Comparisons of EFSA CAG Pesticides and CTD Data

We conducted a proof-of-concept comparison between the number of affected tissues for pesticides grouped into cumulative assessment groups (CAGs) by EFSA and those identified using CGPD tetramers derived from `CTDbase`. While grouping chemicals based on precise mechanisms of action is ideal, EFSA’s Scientific Panel on Plant Protection Products and their Residues (PPR Panel) has acknowledged that grouping can also rely solely on common target organ or system toxicity when detailed mechanistic data is unavailable (*EFSA*, 2008; Committee et al., 2021). Although this approach facilitates risk assessment in data-limited scenarios, it introduces significant uncertainty when evaluating cumulative effects in broadly defined groups (Semino-Beninel et al., 2024). Our comparison evaluates whether CGPD tetramers can effectively capture tissue-specific molecular interactions consistent with EFSA’s CAGs while also highlighting general differences in data availability that may influence grouping outcomes.

When combining the number of pesticides under filtering strategy v5 for human, mouse, and rat data, we identified 63 pesticides overlapping with the EFSA CAG definitions. Figure 5 A shows the number of assigned tissues per pesticide based on `CTD`-derived tetramers and the EFSA CAG reports. For most of the chemicals, especially those with only a few affected tissues in the CAG reports, we detected identical or comparable number of tissues.

**Fig. 5:**
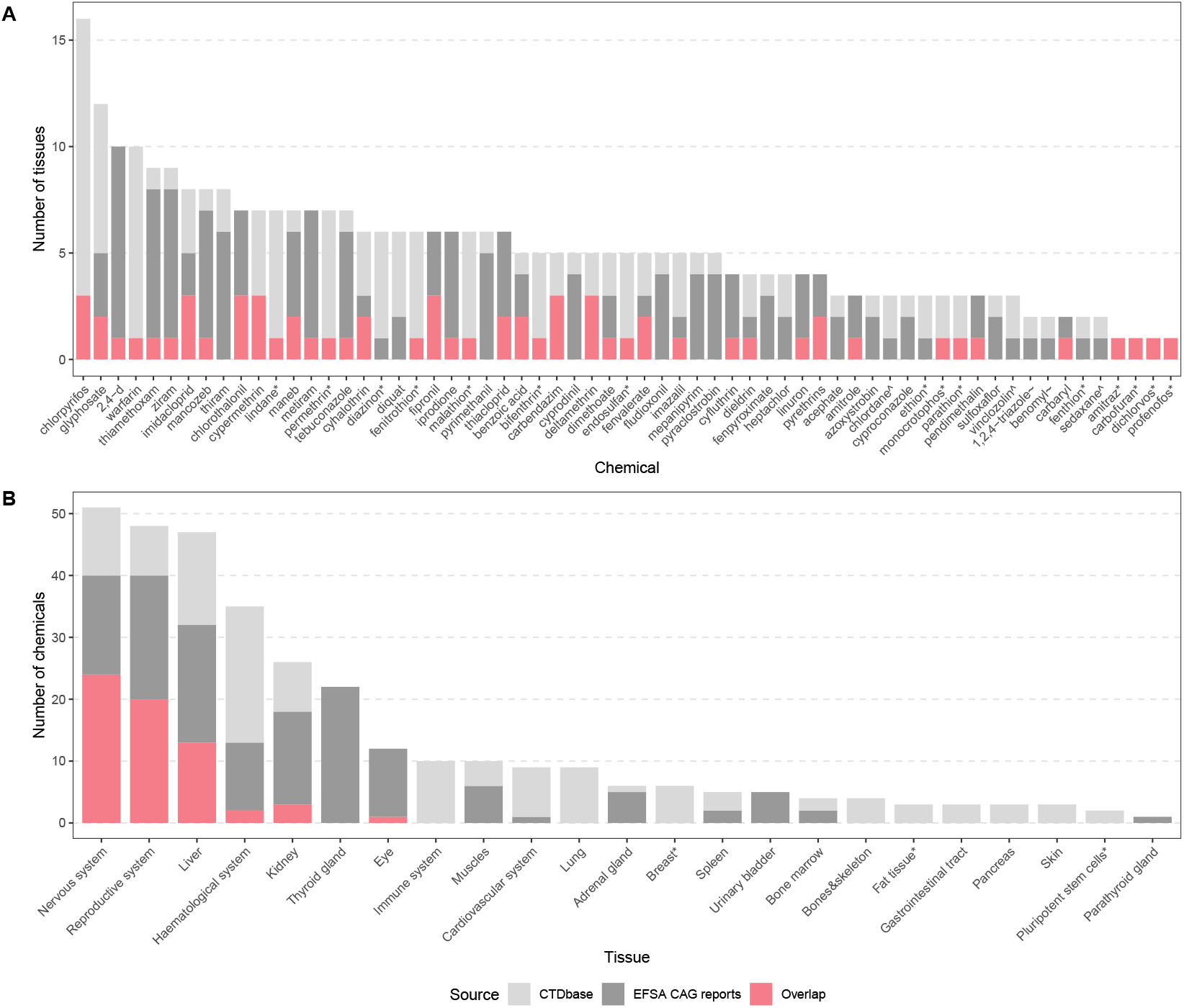
Comparison pesticides and associated tissues between EFSA CAG reports and CTDbase. (A) Comparing 63 pesticides and associated tissues. Pesticides were collected from EFSA CAG reports. `CTD` data was gathered using the most relaxed filtering strategy v5, results for the three target organisms were combined. * denotes 15 pesticides that were exclusively described in the CAG report of the nervous system (EFSA et al., 2019b). ^ marks three pesticides exclusively retrieved from the thyroid gland CAG report (EFSA et al., 2019a). ~ marks two pesticides exclusively retrieved from the reproductive system CAG report (EFSA et al., 2022a). (B) Comparing tissue groups and the number of associated pesticides. * denotes tissue groups that were not covered in the EFSA CAG reports. For both subfigures, ligh grey illustrates `CTD`-based associations, dark grey illustrates associations derived from the EFSA CAG reports, while red highlights associations that were derived from both sources.

A total of 21 pesticides were linked to one tissue in the EFSA CAG reports. Of these, 15 were associated with the nervous system CAG (EFSA et al., 2019b) (marked with *), two with the thyroid gland (EFSA et al., 2019a) (marked with ^), and three with the reproductive system (EFSA et al., 2022a) (marked with ~). For 12 of the 15 nervous system-specific pesticides, we also found molecular interactions with this tissue using `CTD` data. However, no links to the thyroid or reproductive system were found for the remaining pesticides. The absence of associations with the reproductive system-specific pesticides could be due to the particular scope of the CAG report, which focused on transgenerational craniofacial alterations, a feature that is not reliably captured in `CTDbase`. For instance, 1,2,4-triazole was not linked to any reproductive system-associated disease in `CTD` with direct evidence, and benomyl, although linked to prenatal injuries, was annotated only in zebrafish and excluded from our calculations.

In some cases, our analysis revealed significant discrepancies between the number of tissues reported in the EFSA CAG reports and those detected using `CTD` data. For example, pesticides such as 2,4-Dichlorophenoxyacetic acid (EFSA CAGs: 10 tissues), thiamethoxam (8), ziram (8), and metiram (7) were associated with only one or two tissues in the `CTDbase`. Conversely, chlorpyrifos was linked to 16 tissues in `CTDbase`, compared to just three (eye, nervous system, and reproductive system) in the EFSA CAG reports. Similarly, glyphosate and warfarin were associated with 9 and 10 tissues, respectively, in CGPD tetramers, compared to 5 and 1 tissue in the EFSA CAGs. To ensure that these findings were not artifacts caused by relaxed filtering strategies, we analyzed the six pesticides with the most significant discrepancies in their impacted tissues across filtering strategies v1 to v5 (see Supplementary Figure S5). The transition from v1 to v5 resulted in only a slight increase in the number of tissue associations. Notably, even the most stringent filtering strategy (v1) revealed discrepancies, likely due to updated knowledge captured in curated disease interactions from publications post-2012, which were unavailable during the compilation of the initial EFSA CAG report (Nielsen et al., 2012).

In Figure 5 B, we compared pesticide-tissue associations derived `CTD` data with those listed in the EFSA CAG reports. For tissues commonly affected by a broad range of compounds, several additional pesticides were identified with our approach that were not linked to that tissue in the CAG reports.

For the nervous system, 35 pesticides were associated using `CTD` data compared to 40 in the CAG reports, with 24 overlapping chemicals. Eleven pesticides were newly linked to the nervous system based on `CTD`-curated evidence but were not listed as neurotoxic in the CAG report, despite being mentioned in other tissue groups (details in Supplementary Table S5). For instance, azoxystrobin was found to impair neuronal viability and neurite outgrowth in mice (Kang et al., 2021), highlighting potential neurodevelopmental risks. The EFSA CAG report listed 32 liver associated pesticides while our approach linked 28 including 13 overlapping compounds. We identified 15 potentially liver-toxic chemicals with the CGPD tetramers approach being potential candidates for a liver CAG refinement, see Supplementary Table S6. For the reproductive system, eight pesticides were newly associated based on `CTD`-data without being mentioned in the CAG reports (see Supplementary Table S7). The most notable discrepancy was observed for the hematological system, where 24 pesticides were identified using `CTD` data compared to only 13 in the CAG reports, with an overlap of just two (pyrethrins and chlorothalonil; see Supplementary Table S8). Conversely, no pesticides were identified for the thyroid gland using `CTD` data, despite 22 being annotated in the EFSA CAG reports.

These results demonstrate that `CTD`-derived data effectively associates pesticides with established CAG tissues while also uncovering potential novel candidates.

## 4 Cluster Characterization

We explored two use cases to illustrate the potential of our method to identify coherent chemical groups. The vast array of chemical groups generated by our parameter tuning efforts posed a significant challenge, often resulting in overlapping groups across different parameter settings.

To address this complexity, we implemented two scoring procedures: (1) an over-representation analysis focusing on pesticides to identify chemical groups with overlaps to established Cumulative Assessment Groups (CAGs), and (2) an overrepresentation analysis based on the assigned gene lists of each group, using Gene Ontology (GO) terms to uncover insights into potential common modes of action. In the following, we highlight one example of each approach to showcase the versatility and effectiveness of these methods.

### 4.1 Chemical Cluster Linked to Oligospermia

Through an over-representation analysis focused on EFSA CAGs and restricted to pesticides, we identified a noteworthy cluster associated with the reproductive system and the disease oligospermia (MeSH: D009845) in the rat data set, as illustrated in Figure 6. This cluster is associated with seven genes and 19 distinct phenotypes, with bar plots showing the number of unique chemical-gene and chemical-phenotype interactions for these genes and the top 7 phenotypes in Figure 6.

**Fig. 6:**
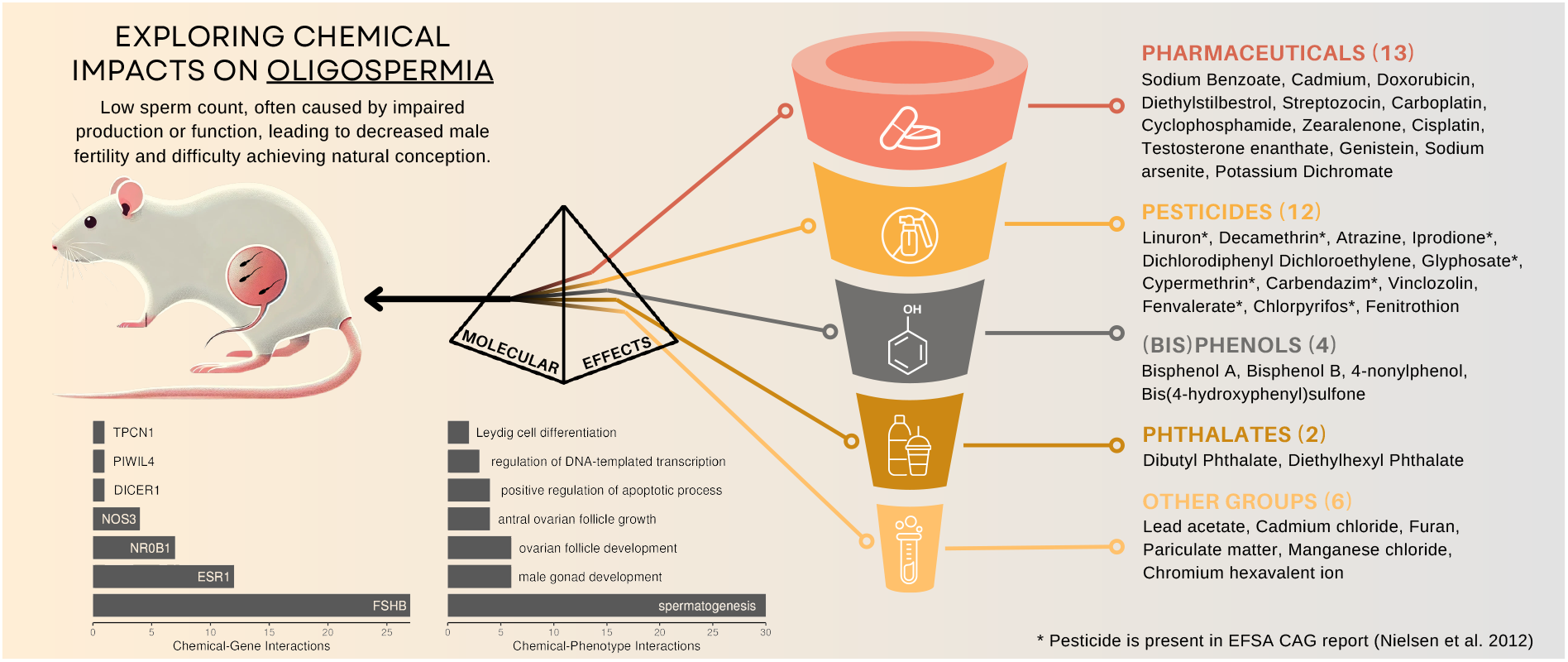
Visualizing chemical cluster linked to Oligospermia. This cluster comprises 37 chemicals associated with the disease Oligospermia in *Rattus norvegicus*, connected through 19 phenotypes and seven genes. Among the 12 pesticides included in this cluster, eight were previously listed in the CAG report by Nielsen et al. (2012) (marked with *) and grouped under CAG 3a ‘Anti-androgenic mode of action’ in the reproductive system. The bottom-left bar plot shows unique chemical-gene interactions for all seven associated genes, while the bottom-right bar plot highlights the top 7 chemical-phenotype interactions. A complete list of associated phenotypes is provided in Supplementary Table S9. Additional details are provided in the text.

The cluster’s main phenotypes included spermatogenesis (GO:0007283), male gonad development (GO:0008584), and ovarian follicle development (GO:0001541), with 30, 6, and 6 unique chemical-phenotype interactions, respectively. Unfortunetaly, `CTDbase` does not keep track of sex-related information in those interactions relevant for CGPD tetramer construction. Therefore, it might happen that female terms are linked with male-specific outcomes and *vice versa*. A complete list of associated phenotypes is provided in Supplementary Table S9.

Associated genes are *Fshb* (chemical-gene interactions: 27), *Esr1* (12), *Nr0b1* (7), *Nos3* (4), *Dicer1* (1), *Piwil4* (1), and *Tpcn1* (1). The genes *Fshb* and *Esr1* are critical regulators of reproductive function, and disruptions in their expression can lead to significant reproductive impairments. For instance, the loss of membrane-associated *Esr1* has been shown to impair sperm maturation in male mice and resulting in reduced fertility (Nanjappa et al., 2016). Similarly, *Fshb* deficiency in rodents leads to a drastic reduction in germ cells, emphasizing its essential role in supporting spermatogenesis and overall reproductive health (Oduwole et al., 2018).

Originally, the cluster grouped 37 chemicals, including eight pesticides listed in the EFSA CAG report (Nielsen et al., 2012) (marked with * in Figure 6). Six of these eight pesticides were included in the CAG Level 4 entitled ‘Compounds with *in vivo* effects that may be attributed to antiandrogenicity and for which inhibition of steroid synthesis is seen *in vitro*’. The remaining two pesticides, carbendazim (C006698) and decamethrin (C017180), had also annotated chemical-gene interactions with key steroid biosynthesis genes such as *Esr1, Fshb, Ar*, and *Star* in `CTD`. All eight pesticides were grouped in CAG 3a entitled “Anti-androgenic mode of action” which was furthermore recommended for cumulative risk assessment by Nielsen et al. (2012).

Four additional pesticides not mentioned in the original EFSA CAG report but listed in the EFSA PARAM catalogue were linked to this cluster: atrazine (D001280), dichlorodiphenyl dichloroethylene (D003633), fenitrothion (D005278), and vinclozolin (C025643). These pesticides were also found to be associated with steroid hormone synthesis genes such as *Fshb* and *Esr1* in our study and, thus, they might be considered as potential CAG refinement candidates. Exposure to atrazine, for example, has been shown to alter the secretion of gonadotropins, including FSH (Wirbisky and Freeman, 2015) leading to decreased sperm count and increased sperm abnormalities in rats as well as human (Abarikwu et al., 2010; Barouki et al., 2012)

The cluster also included 25 non-pesticide chemicals, 13 of which were pharmaceuticals. These pharmaceuticals contained antibiotics such as streptozocin (D013311) and doxorubicin (D004317), estrogen derivatives such as diethylstilbestrol (D004054) and zearalenone (D015025), and immunosuppressive agents such as cyclophosphamide (D003520). Cyclophosphamide, for example, is known to increase the risk of premature menopause and infertility in both males and females (Fusco et al., 2021).

Bisphenols A (BPA, C006780), B (BPB, C492482), and S (BPS, C543008) were also grouped in this cluster. BPA, initially developed as a pharmaceutical estrogen replacement, has since been widely used in consumer products. Its reproductive, metabolic, and developmental effects led to its classification as an endocrine-disrupting chemical and a substance of high concern^14^, which eventually led to its ban in products like baby bottles and food packaging. One of BPA’s substitute, BPB, shares strong structural similarity and was shown to have similar endocrine-disrupting effects, such as decreased testosterone production and estrogenic activity (Ullah et al., 2018b,a) A review on the evidence of BPB’s endocrine potentials is given by Serra et al. (2019). A second substitute for BPA is BPS, which has been shown to have comparable estrogenic activity (Grignard et al., 2012), disrupting function of the androgen receptor (Siracusa et al., 2018) and effects on sperm count (Zalmanova et al., 2016).

The cluster also contained two phthalates, diethylhexyl phthalate (DEHP, D004051) and dibutyl phthalate (DBP, D003993), both interacting with genes such as *Fshb, Esr1*, and *Ar*. Both compounds are extensively used as plasticizers in PVC products and are common in household items, cosmetics, and personal care products, although the highest exposure comes through food (Rowdhwal and Chen, 2018). The EU Commision classified these substances as toxic for reproduction in their regulation No 143/2011 on 17 February 2011^15^.

A potential substitute for plasticizers like DEHP is 2,5-furandicarboxylic acid (FDCA). While FDCA is listed in `CTDbase`, it lacks curated interactions and respective CGPD tetramers. However, its precursor, furan (C039281), is included in this cluster and is linked to public health risks identified by EFSA in 2017 (EFSA Panel on Contaminants in the Food Chain (CONTAM) et al., 2017). Furthermore, the European Commision recommended its monitoring in food products in 2022^16^.

This example illustrates the potential of CGPD tetramer-based clustering to provide candidates for risk assessors, facilitate the grouping of compounds for joint/cumulative assessment, and bridge gaps in existing regulatory frameworks by integrating NAM-derived and literature-based data and associations. The identification of shared pathways, such as interactions with steroid hormone synthesis genes, highlights the utility of our method in prioritizing compounds for regulatory evaluation and further investigation of their endocrine-disrupting potential. By grouping chemicals across classes - spanning pesticides, pharmaceuticals, and industrial chemicals - our results emphasize the need to extend risk assessments towards joint considerations of chemicals with diverse applications, which are currently governed by separate regulatory frameworks.

### 4.2 Chemical Cluster Linked to Hyperglycemia

Within the rat hematological system, we identified a distinct cluster of 31 chemicals associated with hyperglycemia, a condition characterized by elevated blood glucose levels. This cluster was linked to the phenotypes glucose homeostasis (GO:0042593) and positive regulation of insulin secretion (GO:0032024). Eight genes were associated with this cluster, all of which play critical roles in glucose metabolism and insulin signaling: *Gck* (Glucokinase), *Il6* (Interleukin 6), *Ins1* (Insulin 1), *Ins2* (Insulin 2), *Insr* (Insulin Receptor), *Lepr* (Leptin Receptor), *Lep* (Leptin), and *Adipoq* (Adiponectin). Dysregulation of these genes has been linked to impaired glucose regulation, insulin resistance, diminished insulin production, and increased risk of type 2 diabetes (Abu Aqel et al., 2024; Kristiansen and Mandrup-Poulsen, 2005; Kadowaki et al., 2006; Yadav et al., 2013; Chandrasekaran and Weiskirchen, 2024).

This cluster was identified based on an overrepresentation analysis of associated genes, resulting in an enrichment of 10 different GO terms with a gene ratio of 1, meaning that all eight genes are listed in those terms. Those terms encompassed glucose homeostasis (GO:0042593) and glucose metabolic process (GO:0006006) but also other carbohydrate-related terms: hexose metabolic process (GO:0019318), monosaccharide metabolic process (GO:0005996), carbohydrate homeostasis (GO:0033500), and carbohydrate metabolic process (GO:0005975), see Supplementary Table S10. The cluster appeared consistently across 40 different parameter settings using the PD-based grouping and filtering strategies v2 to v5 (Supplementary Table S12) containing 31 chemicals of various classes, including six pesticides and 16 pharmaceuticals (details are given in Supplementary Table S11).

Among the six pesticides, acephate (C001969), chlorpyrifos (D004390), and diazinon (D003976) are well-known organophosphates insecticides. Although primarily used as cholinesterase inhibitors targeting the nervous system of insects, these compounds have been associated with hyperglycemia through mechanisms like pancreatic *β*-cell damage, increased hepatic gluconeogenesis, and insulin resistance (Chung et al., 2021; Lasram et al., 2014). For example, acephate induces reversible hyperglycemia in rats, often in co-existence with hypercorticosteronemia (Deotare and Chakrabarti, 1981; Joshi and Rajini, 2009), while chlorpyrifos and diazinon significantly increase blood glucose levels, particularly in female animals (Farkhondeh et al., 2020, 2021).

The neonicotinoid pesticide Imidacloprid (C082359) was designed to mimic the neuroactive effects of nicotine (both found in this cluster), by acting on the nicotinic acetylcholine receptors (nAChRs). While imidacloprid exhibits greater selectivity for insect nAChRs and is considered less toxic to mammals compared to organophosphates, studies have shown that it can disrupt glucose metabolism, leading to adiposity and glucose homeostasis disturbances in rodents (Kim et al., 2013; Sun et al., 2016, 2017). Nicotine itself has proven effects on blood glucose levels by interfering with glucoregulatory hormone release via nAChR activation in pancreatic *β*-cells (Chen et al., 2023).

Additional chemical classes of this cluster included plasticizers, e.g., bisphenol A (BPA; C006780), heavy metals such as lead (D007854) and arsenic (D001151), and brominated flame retardants. BPA has been shown to increase blood glucose levels and induce insulin resistance in a dose-dependent manner (Moghaddam et al., 2015). Lead promotes hyperglycemia through increased hepatic gluconeogenesis (Wan et al., 2021), while arsenic exposure impairs glucose tolerance and increases diabetes risk (Navas-Acien et al., 2006). Brominated flame retardants, such as 2,2’,4,4’-Tetrabromodiphenyl Ether (C511295) and Decabromobiphenyl Ether (C010902), have been associated with disrupted glucose homeostasis and an increased incidence of diabetes (Zhang et al., 2016).

Pharmaceuticals within the cluster include chemotherapy agents, e.g., streptozocin (D013311), which induces diabetes in animal models by selectively targeting pancreatic *β*-cells (Ventura-Sobrevilla et al., 2011), and dexamethasone (D003907), a corticosteroid that elevates blood glucose through hepatic gluconeogenesis, particularly in at-risk individuals (Lukins and Manninen, 2005; Purushothaman et al., 2018). Other drugs such as atorvastatin (D000069059), a cholesterol-lowering statin, and arsenicals like arsenic trioxide (D000077237) and Sodium Arsenite (C017947) were linked to impaired glucose regulation (Ghafghazi et al., 1980; Koh et al., 2010; Reichl et al., 1990).

Conversely, the cluster also included drugs that are primarily prescribed as anti-diabetic medication such as Dapagliflozin (C529054), Glyburide (D005905), and Metformin (D008687) These compounds, while aiming to lower blood glucose, operate via distinct mechanisms: metformin reduces hepatic glucose production and improves insulin sensitivity (Harada, 2020), glyburide stimulates insulin secretion from pancreatic *β*-cells (Qureshi et al., 2018), and dapagliflozin enhances glucose excretion via inhibition of sodium-glucose cotransporter 2 (Sglt2) (Henry et al., 2018).

The hyperglycemia cluster showcases the ability of our methodology to group chemicals with diverse impacts on glucose homeostasis, reflecting both therapeutic and harmful effects. This is primarily due to the application of filtering strategies v2 to v5, where ‘marker/mechanism’ and ‘therapeutic’ disease-relations are included in the calculation of CGPD tetramers.

Under filtering strategy v1, which excludes therapeutic associations, a smaller cluster of 19 chemicals was identified with the same associated disease and phenotypes as for the cluster described above. Six of the eight previously asspciated genes were also linked, solely omitting *Ins1* and *Adipoq*. In this cluster, pesticides and pharmaceuticals linked to negative effects on glucose homeostasis were retained, but compounds with beneficial effects, such as metformin, glyburide, or plant-based compounds like curcumin (D003474) and quercetin (D011794) were excluded. Interestingly, dapagliflozin remained in this cluster despite its known glucose-lowering properties. Notably, hyperglycemic pesticides like acephate and diazinon were absent due to the stricter filtering criteria (Supplementary Table S11). The described differences highlight the influence of parameter settings on cluster composition and demonstrate the ability of our methodology to identify and refine groups of chemicals potentially associated with impairments of biological functions such as glucose regulation. Supplementary Table S12 lists different parameter settings that returned the discussed cluster.

## 5 Conclusions

In this study, we demonstrate the utility of CGPD tetramers derived from `CTDbase` for identifying and grouping chemicals with shared associations to biological and toxicological effects, offering a holistic and powerful approach for chemical grouping and clustering across various chemical classes. The results highlight the potential of our methodology to inform chemical risk assessment and regulatory decision-making while underscoring challenges associated with data availability, data quality, and group identification strategies.

Our study illustrates both the potential and the complexities of leveraging omics-derived data for chemical grouping. Using curated data from `CTDbase`, we grouped chemicals across diverse categories, offering new insights into their cumulative effects and molecular mechanisms. The method’s validation is based on the observed overlap with cumulative assessment groups (CAGs) of pesticides, while the identification of additional chemicals emphasizes its applicability for refining risk assessments.

The quality and coverage of data in `CTDbase` play a critical role in the robustness of our methodology. As a manually curated resource, `CTDbase` relies on published research, which introduces inherent time lags and biases. Legacy chemicals, such as banned pesticides, may be overrepresented compared to newer compounds due to the database’s retrospective nature. This discrepancy can hinder the effectiveness of immediately including newly approved chemicals, for example, in existing CAGs. The quality and completeness of data in `CTDbase` are substantially influenced by the manual curation process, which, while thorough, can lead to (1) loss of information due to incomplete curation of available data and (2) the incorporation of potentially false interactions. For example, a transcriptomics study on clothianidin by Alarcan et al. (2021) identified 2986 differentially expressed genes using RNA-Seq in rats. However, `CTDbase` annotated only ten genes manually selected and validated by qRT-PCR, representing only a fraction of the overall dataset. Based on a follow-up paper using the same data set, `CTD` lists 22 different chemical-gene interactions. However, those genes were used as target genes for RT-PCR, and, in fact, only three of them were differentially expressed (Sprenger et al., 2022). While the sheer volume of `CTDbase` data helps to mitigate some noise, such inconsistencies emphasize the need for cautious interpretation and continuous refinement of curation methodologies.

The vast number of chemical groups generated through our evaluation procedure, which utilized multiple filtering strategies and parameter combinations, reflects both the flexibility and complexity of our approach. To manage this complexity and demonstrate its reliability, we developed two complementary methods to identify meaningful clusters: an overrepresentation analysis against pesticide CAGs and a GO-based enrichment analysis of gene lists. However, selecting relevant clusters ultimately depends on the specific research or regulatory question, emphasizing the need for tailored evaluation procedures.

Our approach extends beyond pesticides, encompassing pharmaceuticals and industrial chemicals of regulatory importance. This holistic perspective is pivotal in uncovering the cumulative impacts of diverse chemical exposures on human health and the environment. By identifying shared effects on genes, phenotypes, and diseases, our methodology aligns with the foundational work on pesticide CAGs by Nielsen et al. (2012) and highlights the multifaceted nature of chemical risks. It advocates for a comprehensive assessment framework that can better support regulatory decisions and public health policies.

Despite these advancements, the direction, quantity, and effect size of a particular response, which is critical for understanding chemical impact, remains unrecognized in our current grouping strategy. This limitation complicates the interpretation of the results. Incorporating directional data for chemical-gene relations, similar to our existing handling of evidence types in disease interactions, could significantly refine future analyses. Moving forward, integrating directional effects, improved tissue-specific annotations, and expanded organism combinations will enhance the accuracy and interpretability of our results.

Ultimately, this methodology marks a significant step toward more comprehensive, data-driven approaches that can support chemical risk assessments. By bridging gaps across diverse regulatory frameworks being needed for distinct chemical classes, we address the challenges posed by cumulative exposures and contribute to advancing risk assessment practices in alignment with emerging regulatory goals.

## Acknowledgements

The authors are grateful to Giovanni Iacono and Sara Levorato from EFSA for fruitful discussions.

## Supplementary Material

The Supplementary Material is available at Zenodo with the DOI https://doi.org/10.5281/zenodo.14724322. A pdf file containing Supplementary Tables and Figures is also part of this repository and individually accessible with this link: https://zenodo.org/records/14724322/files/Canzler_et_al_Grouping_Supplementary_Material.pdf

## Footnotes

1 https://www.fda.gov/media/182478/download

2 https://echa.europa.eu/regulations/reach/understanding-reach

3 https://ctdbase.org/

4 https://food.ec.europa.eu/horizontal-topics/farm-fork-strategy_en

5 https://environment.ec.europa.eu/strategy/chemicals-strategy_en

6 https://pubchem.ncbi.nlm.nih.gov/idexchange/idexchange.cgi

7 https://zenodo.org/records/14724322/files/2023_03_20_ctd_pubchem_sqlite.db.gz

8 https://ftp.ncbi.nlm.nih.gov/gene/DATA/gene_orthologs.gz

9 https://ftp.ncbi.nlm.nih.gov/gene/DATA/gene2go.gz

10 https://zenodo.org/records/14724322/files/Chemical_organism_associations.tsv

11 https://codebase.helmholtz.cloud/department-computational-biology/efsa_cags/chemical_grouping_workflow

12 https://doi.org/10.5281/zenodo.7590216

13 https://go.drugbank.com/

14 https://echa.europa.eu/documents/10162/ac9efb97-c06b-d1a7-2823-5dc69208a238

15 http://data.europa.eu/eli/reg/2011/143/oj

16 http://data.europa.eu/eli/reco/2022/495/oj

