## Supplementary Material for "From Toxicogenomics Data to Cumulative Assessment Groups: A Mechanistic Framework for Chemical Grouping"

**Noname manuscript No.**
(will be inserted by the editor)

### Supplementary Material

#### From Toxicogenomics Data to Cumulative Assessment Groups: A Mechanistic Framework for Chemical Grouping

**Sebastian Canzler · Julienne Lehmann · Jana Schor · Wibke Busch · Jörg Hackermüller**

Received: date / Accepted: date

S.Canzler · J.Lehmann · J.Schor · J.Hackermüller
Department of Computational Biology and Chemistry, Helmholtz Centre for Environmental Research - UFZ, 04318 Leipzig, Germany


W. Busch
Department Ecotoxicology Helmholtz Centre for Environmental Research - UFZ, 04318 Leipzig, Germany

J. Lehmann
HIV Cure Research Center, Department of Internal Medicine and Pediatrics, Ghent University Hospital, Ghent University, 9000 Ghent, Belgium

J. Hackermüller
Department of Computer Science, Leipzig University, 04109 Leipzig, Germany

Table S1: Summary of source files that have been used to gather CTD and PubChem data.

| Source | Property | File URL |
|---|---|---|
| CTD | Chem - Gene | https://ctdbase.org/reports/CTD_chem_gene_ixns.tsv.gz |
| CTD | Chem - Gene Types | https://ctdbase.org/reports/CTD_chem_gene_ixn_types.tsv |
| CTD | Chem - Disease | https://ctdbase.org/reports/CTD_chemicals_diseases.tsv.gz |
| CTD | Chem - Pathway | https://ctdbase.org/reports/CTD_chem_pathways_enriched.tsv.gz |
| CTD | Chem - GO | https://ctdbase.org/reports/CTD_chem_go_enriched.tsv.gz |
| CTD | Gene - Pathway | https://ctdbase.org/reports/CTD_genes_pathways.tsv.gz |
| CTD | Disease - Pathway | https://ctdbase.org/reports/CTD_diseases_pathways.tsv.gz |
| CTD | Chemicals | https://ctdbase.org/reports/CTD_chemicals.tsv.gz |
| CTD | Diseases | https://ctdbase.org/reports/CTD_diseases.tsv.gz |
| CTD | Genes | https://ctdbase.org/reports/CTD_genes.tsv.gz |
| CTD | Pathways | https://ctdbase.org/reports/CTD_pathways.tsv.gz |
| CTD | Anatomy terms | https://ctdbase.org/reports/CTD_anatomy.tsv.gz |
| CTD | Exposure | https://ctdbase.org/reports/CTD_exposure_events.tsv.gz |
| CTD | Exp. studies | https://ctdbase.org/reports/CTD_exposure_studies.tsv.gz |
| CTD | Chem - Phenotype | https://ctdbase.org/reports/CTD_pheno_term_ixns.tsv.gz |
| CTD | Gene - Disease | https://ctdbase.org/reports/CTD_genes_diseases.tsv.gz |
| PubChem | InChI key | https://ftp.ncbi.nlm.nih.gov/pubchem/Compound/Extras/CID-InChI-Key.gz |
| PubChem | SMILES | https://ftp.ncbi.nlm.nih.gov/pubchem/Compound/Extras/CID-SMILES.gz |
| PubChem | Molec. mass, Formula | https://ftp.ncbi.nlm.nih.gov/pubchem/Compound/Extras/CID-Mass.gz |
| PubChem | Title | https://ftp.ncbi.nlm.nih.gov/pubchem/Compound/Extras/CID-Title.gz |
| PubChem | CAS, ChEBI DTXSID | https://ftp.ncbi.nlm.nih.gov/pubchem/Compound/Extras/CID-Synonym-filtered.gz |

Table S2: Table summarizing the SQLite structure. The data in the tables starting with 'CID' was collected from PubChem and originated from different source files. Tables starting with 'CTD' originated from CTD. Table names are equivalent to the CTD file name. The snapshot date of the data set was March 20th, 2023.

| Table name | # Columns | # Entries |
|---|---|---|
| CID_chemicalproperties | 6 | 12041 |
| CID_IDs | 6 | 16332 |
| CTD_anatomy | 9 | 1844 |
| CTD_chem_gene_ixns | 12 | 2487743 |
| CTD_chem_gene_ixn_types | 4 | 53 |
| CTD_chem_go_enriched | 13 | 6237303 |
| CTD_chemicals | 8 | 175847 |
| CTD_chemicals_diseases | 10 | 8276711 |
| CTD_chem_pathways_enriched | 11 | 1483203 |
| CTD_diseases | 9 | 13270 |
| CTD_diseases_pathways | 5 | 609539 |
| CTD_exposure_events | 43 | 204735 |
| CTD_exposure_studies | 10 | 3304 |
| CTD_genes | 8 | 589897 |
| CTD_genes_diseases | 9 | 100715108 |
| CTD_genes_pathways | 4 | 135783 |
| CTD_pathways | 2 | 2567 |
| CTD_pheno_term_ixns | 13 | 340861 |

#### S0.1 Tissue groups

**Adrenal gland**

- Adrenal Cortex
- Adrenal Glands

**Bone marrow**

- Mesenchymal Stem Cells
- Myeloid Cells
- Bone Marrow
- Endothelial Progenitor Cells
- Cancellous Bone
- Bone Marrow Cells

**Bones/skeleton**

- Osteoblasts
- Skull
- Synovial Membrane
- Bone and Bones
- Jaw
- Maxilla
- Orbit
- Turbinates
- Nucleus Pulposus
- Lumbar Vertebrae
- Cartilage
- Articular
- Ankle Joint
- Joint Capsule
- Knee Joint
- Osteocytes
- Femur
- Leg Bones
- Tibia
- Spine
- Joints
- Cartilage, Articular
- Menisci, Tibial
- Skeleton
- Growth Plate
- Mandible
- Tarsal Joints
- Periosteum

**Cardiovascular system**

- Cardiovascular System
- Blood Vessels
- Exudates and Transudates
- Arteries
- Aorta
- Brachial Artery
- Middle Cerebral Artery
- Coronary Vessels
- Coronary Sinus
- Umbilical Arteries
- Heart
- Heart Atria
- Heart Ventricles
- Myocardium
- Myocytes, Cardiac
- Endothelial Cells
- Human Umbilical Vein Endothelial Cells
- Mesenteric Arteries
- Pulmonary Artery
- Microvessels
- Endothelium, Vascular
- Veins
- Umbilical Veins
- Aortic Valve
- Myoblasts, Cardiac
- Blood-Brain Barrier
- Aorta, Abdominal
- Aorta, Thoracic
- Sinus of Valsalva
- Brachiocephalic Trunk
- Carotid Arteries
- Femoral Artery
- Mesenteric Artery, Superior
- Uterine Artery
- Venules
- Cerebral Veins
- Hepatic Veins
- Portal Vein
- Ductus Arteriosus
- Pericardium
- Mitochondria, Heart
- Blood-Testis Barrier
- Adventitia
- Arterioles
- Basilar Artery
- Cerebral Arteries
- Anterior Cerebral Artery
- Circle of Willis
- Meningeal Arteries
- Thoracic Arteries
- Capillaries
- Retinal Vessels
- Tunica Intima
- Portal System
- Mesenteric Veins

**Eye**

- Eye
- Retinal Pigment Epithelium
- Conjunctiva
- Cornea
- Epithelium, Corneal
- Lens, Crystalline
- Trabecular Meshwork
- Retina
- Photoreceptor Cells
- Retinal Ganglion Cells
- Tenon Capsule
- Uvea
- Iris
- Lacrimal Apparatus
- Photoreceptor Cells, Vertebrate
- Retinal Neurons
- Retinal Horizontal Cells
- Ciliary Body
- Corneal Stroma

**Gallbladder**

- Gallbladder

**Haematological system**

- Granulocyte Precursor Cells
- Lymphoid Progenitor Cells
- Myeloid Progenitor Cells
- Granulocyte-Macrophage Progenitor Cells
- U937 Cells
- Erythroid Precursor Cells
- Blood
- Blood Cells
- Serum
- Blood Buffy Coat
- Plasma
- Erythrocytes
- Blood Platelets
- Fetal Blood
- Erythrocyte Membrane
- Hematopoietic Stem Cells
- Precursor Cells, B-Lymphoid
- Precursor Cells, T-Lymphoid
- Erythroblasts
- Megakaryocyte Progenitor Cells
- Megakaryocytes
- Reticulocytes
- Erythroid Cells
- Thymocytes

**Kidney**

- Madin Darby Canine Kidney Cells
- Kidney
- Kidney Glomerulus
- Mesangial Cells
- Kidney Medulla
- Kidney Tubules, Proximal
- Kidney Cortex
- Podocytes
- Kidney Tubules
- Juxtaglomerular Apparatus
- Kidney Tubules, Collecting
- Kidney Tubules, Distal

**Liver**

- Hep G2 Cells
- Microsomes, Liver
- Hepatic Stellate Cells
- Hepatocytes
- Bile Ducts
- Liver
- Bile Ducts, Intrahepatic
- Bile Canaliculi
- Mitochondria, Liver
- Bile

**Muscles**

- Muscle, Skeletal
- Muscle, Smooth, Vascular
- Muscles
- Muscle, Striated
- Satellite Cells, Skeletal Muscle
- Muscle Cells
- Myoblasts, Skeletal
- Myocytes, Smooth Muscle
- Rectus Abdominis
- Quadriceps Muscle
- Diaphragm
- Muscle Fibers, Skeletal
- Myoblasts
- Abdominal Muscles
- Tendons
- Achilles Tendon

**Nervous system**

- Nervous System
- Corticotrophs
- PC12 Cells
- Brain
- Substantia Nigra
- Ventral Tegmental Area
- Hippocampus
- CA1 Region, Hippocampal
- Parahippocampal Gyrus
- Entorhinal Cortex
- Corpus Striatum
- Nucleus Accumbens
- Cerebral Cortex
- Frontal Lobe
- Neocortex
- Spinal Cord
- Astrocytes
- Oligodendroglia
- Neurons
- Dopaminergic Neurons
- Neural Stem Cells
- Mesencephalon
- Cerebellum
- CA2 Region, Hippocampal
- CA3 Region, Hippocampal
- Dentate Gyrus
- Somatotrophs
- Cerebrum
- Meninges
- Neuroglia
- Ependymoglial Cells
- Microglia
- Schwann Cells
- Peripheral Nervous System
- Sympathetic Nervous System
- Neuroepithelial Cells
- Central Nervous System
- Brain Stem
- Cerebral Peduncle
- Pars Compacta
- Tegmentum Mesencephali
- Locus Coeruleus
- Inferior Colliculi
- Superior Colliculi
- Cerebellar Cortex
- Purkinje Cells
- Pons
- Facial Nucleus
- Superior Olivary Complex
- Medulla Oblongata
- Cerebral Ventricles
- Lateral Ventricles
- Limbic System
- Amygdala
- Pineal Gland
- Fornix, Brain
- Hypothalamus
- Paraventricular Hypothalamic Nucleus
- Arcuate Nucleus of Hypothalamus
- Median Eminence
- Gonadotrophs
- Thalamus
- Telencephalon
- Globus Pallidus
- Neostriatum
- Putamen
- Motor Cortex
- Prefrontal Cortex
- Somatosensory Cortex
- Basal Forebrain
- Olfactory Bulb
- Septal Nuclei
- Corpus Callosum
- White Matter
- Pyramidal Tracts
- Spinal Cord Dorsal Horn
- Spinal Cord Ventral Horn
- Ganglia, Spinal
- Olfactory Pathways
- Medial Forebrain Bundle
- Oligodendrocyte Precursor Cells
- Cholinergic Neurons
- Neurites
- GABAergic Neurons
- Hair Cells, Auditory
- Sensory Receptor Cells
- Motor Neurons
- Pyramidal Cells
- Synapses
- Anterior Horn Cells
- Hair Cells, Auditory, Inner
- Hair Cells, Auditory, Outer
- Olfactory Receptor Neurons
- Myenteric Plexus
- Optic Nerve
- Sciatic Nerve
- Neuroendocrine Cells
- Raphe Nuclei
- Dorsal Raphe Nucleus
- Trigeminal Nucleus, Spinal
- Olivary Nucleus
- Solitary Nucleus
- Choroid Plexus
- Basolateral Nuclear Complex
- Central Amygdaloid Nucleus
- Corticomedial Nuclear Complex
- Preoptic Area
- Supraoptic Nucleus
- Gyrus Cinguli
- Prosencephalon
- Geniculate Bodies
- Lateral Thalamic Nuclei
- Midline Thalamic Nuclei
- Basal Ganglia
- Parietal Lobe
- Sensorimotor Cortex
- Piriform Cortex
- Temporal Lobe
- Posterior Horn Cells
- Dura Mater
- Pia Mater
- Cervical Cord
- Spiral Ganglion
- Myelin Sheath
- Dendritic Spines
- Autonomic Pathways
- Peripheral Nerves
- Spinal Nerves
- Stellate Ganglion
- Superior Cervical Ganglion
- Carotid Body
- Vagus Nerve
- Spinal Nerve Roots

**Parathyroid gland**

- Parathyroid Glands

**Reproductive system**

- Semen
- Placenta
- Endometrium
- Trophoblasts
- Spermatozoa
- Ovary
- Granulosa Cells
- Testis
- Prostate
- PC-3 Cells
- Genitalia, Male
- Penis
- Genitalia, Female
- Oocytes
- Follicular Fluid
- Luteal Cells
- Cervix Uteri
- Ovarian Follicle
- Foreskin
- Mesoderm
- Uterus
- Germ Cells
- Sertoli Cells
- Chorionic Villi
- Cumulus Cells
- Decidua
- Leydig Cells
- Spermatocytes
- Epididymis
- Seminiferous Tubules
- Spermatogonia
- Seminal Vesicles
- Theca Cells
- Gonads
- Zygote
- Vagina
- Spermatids
- Corpus Luteum
- Seminiferous Epithelium
- Sperm Head
- Rete Testis
- Vas Deferens
- Fallopian Tubes
- Urethra
- Genitalia

**Breast**

- Mammary Glands
- Mammary Glands, Human
- Mammary Glands, Animal
- MCF-7 Cells
- Breast

**Spleen**

- Spleen

**Thyroid gland**

- Thyroid Epithelial Cells
- Thyroid Gland

**Urinary bladder**

- Urothelium
- Urinary Bladder
- Urinary Tract
- Ureter
- Urothelium

**Gastrointestinal tract**

- Gastrointestinal Tract
- Colon
- Intestine, Small
- Duodenum
- Mouth
- Taste Buds
- Stomach
- Dental Pulp
- Caco-2 Cells
- HCT116 Cells
- Intestines
- Intestine, Large
- Cecum
- Rectum
- Submandibular Gland
- Tongue
- Pharynx
- Esophagus
- Esophageal Mucosa
- Gastric Mucosa
- Periodontium
- Gingiva
- Periodontal Ligament
- Tooth
- Bicuspid
- Molar
- Dental Papilla
- Hypopharynx
- Nasopharynx
- Peritoneal Cavity
- Mouth Floor
- Omentum
- Intestinal Mucosa
- Ileum
- Jejunum
- Lower Gastrointestinal Tract
- Upper Gastrointestinal Tract
- Dental Cementum
- Enamel Organ
- Enteroendocrine Cells
- Enterocytes
- Incisor
- Dental Enamel
- Parietal Cells, Gastric
- HT29 Cells
- Interstitial Cells of Cajal

**Immune system**

- RAW 264.7 Cells
- Osteoclasts
- Synoviocytes
- Immune System
- B-Lymphocytes
- B-Lymphocyte Subsets
- Plasma Cells
- Dendritic Cells
- Langerhans Cells
- Leukocytes
- Granulocytes
- HL-60 Cells
- Basophils
- Eosinophils
- Neutrophils
- Leukocytes, Mononuclear
- Lymphocytes
- Monocytes
- Endothelium, Lymphatic
- Mast Cells
- Macrophages
- Foam Cells
- Killer Cells, Natural
- T-Lymphocytes, Helper-Inducer
- T-Lymphocytes, Regulatory
- T-Lymphocytes
- CD4-Positive T-Lymphocytes
- CD8-Positive T-Lymphocytes
- Jurkat Cells
- Th17 Cells
- Lymphatic System
- Lymphoid Tissue
- Histiocytes
- Th1 Cells
- Th2 Cells
- Kupffer Cells
- Macrophages, Alveolar
- Macrophages, Peritoneal
- Phagocytes
- Myeloid-Derived Suppressor Cells
- Lymph
- THP-1 Cells

**Lung**

- A549 Cells
- Sputum
- Lung
- Bronchi
- Alveolar Epithelial Cells
- Bronchioles
- Pulmonary Alveoli
- Pleura
- Trachea
- Respiratory System

**Lymph Nodes**

- Lymph Nodes
- Germinal Center

**Pancreas**

- Islets of Langerhans
- Insulin-Secreting Cells
- Pancreatic Ducts
- Glucagon-Secreting Cells
- Pancreas
- Pancreas, Exocrine
- Pancreatic Stellate Cells

**Pituitary gland**

- Pituitary Gland
- Pituitary Gland, Anterior

**Salivary gland**

- Saliva
- Salivary Glands
- Parotid Gland

**Skin**

- HaCaT Cells
- Keratinocytes
- Melanocytes
- Skin
- Dermis
- Epidermis
- Hair Follicle
- Sebaceous Glands
- Epidermal Cells

**Thymus**

- Thymus Gland

**Fat tissues**

- Adipocytes
- Adipocytes, White
- Adipocytes, Brown
- Adipose Tissue
- Adipose Tissue, White
- Adipose Tissue, Brown
- Subcutaneous Fat
- Subcutaneous Tissue
- Intra-Abdominal Fat
- 3T3-L1 Cells

**Embrionic stem cells**

- HEK293 Cells
- Embryonic Stem Cells, Cultured
- Human Embryonic Stem Cells
- Embryonic Structures
- Mouse Embryonic Stem Cells
- Embryonic Germ Cells
- Embryonal Carcinoma Stem Cells

Table S3: Pesticide-related MeSH terms that were used to identify pesticides in CTD.

| MeSH Term | Number of pesticides in CTD |
|---|---|
| Insecticides | 151 |
| Herbicides | 78 |
| Fungicides | 63 |
| Pesticides | 19 |
| Rodenticides | 10 |
| Insect Repellents | 9 |
| Chemosterilants | 5 |
| Molluscacides | 3 |
| Defoliants | 3 |
| Acaricides | 2 |
| Pesticide Synergists | 1 |

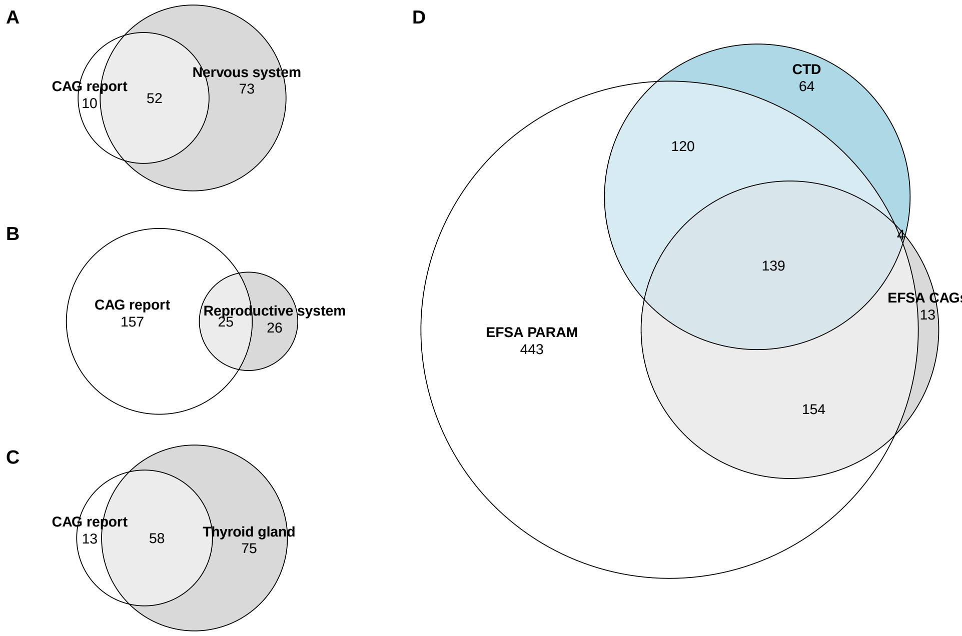


Fig. S1: Overlap of pesticides collected from different sources. Subfigures A-C focus on tissue-specific overlaps between CAGs initially proposed by Nielsen et al. (2012) and those subsequently established by EFSA [2019,2019a,2022]. Pesticides collected from the original CAG report in 2012 are denoted as 'CAG report' in those subfigures. (A) Pesticides impacting the nervous system. (B) Pesticides impacting the reproductive system and the updated report focusing on the craniofacial alterations. (C) Pesticides impacting the thyroid gland. (D) Overlap of pesticides collected from the EFSA PARAM catalogue, the combined CAG reports, and pesticides collected from CTDbase using the MeSH dictionary.

Table S4: MeSH terms that were listed in the MeSH Browser tree view as children of "Physiological Effects of Drugs [D27.505.696]", which are associated with pharmaceuticals.

| MeSH Term | MeSH Term |
|---|---|
| Physiological Effects of Drugs | Abuse-Deterrent Formulations |
| Aversive Agents | Antipyretics |
| Antispermatogenic Agents | Sperm Immobilizing Agents |
| Spermatocidal Agents | Spermatogenesis-Blocking Agents |
| Astringents | Bone Density Conservation Agents |
| Calcium-Regulating Hormones and Agents | Calcium Channel Agonists |
| Calcium Channel Blockers | Central Nervous System Depressants |
| Anesthetics | Anesthetics, Combined |
| Anesthetics, General | Anesthetics, Inhalation |
| Anesthetics, Intravenous | Anesthetics, Dissociative |
| Anesthetics, Local | Hypnotics and Sedatives |
| Sleep Aids, Pharmaceutical | Orexin Receptor Antagonists |
| Narcotics | Analgesics, Opioid |
| Tranquilizing Agents | Anti-Anxiety Agents |
| Antimanic Agents | Antipsychotic Agents |
| Central Nervous System Stimulants | Aphrodisiacs |
| Appetite Stimulants | Convulsants |
| Cerumenolytic Agents | Emetics |
| Endocrine Disruptors | Galactogogues |
| Growth Substances | Angiogenesis Modulating Agents |
| Angiogenesis Inducing Agents | Angiogenesis Inhibitors |
| Growth Inhibitors | Angiogenesis Inhibitors |
| Plant Growth Regulators | Hallucinogens |
| Hormones, Hormone Substitutes, and Hormone Antagonists | Hormone Antagonists |
| Androgen Antagonists | Androgen Receptor Antagonists |
| Nonsteroidal Anti-Androgens | Antithyroid Agents |
| Calcimimetic Agents | Estrogen Antagonists |
| Aromatase Inhibitors | Estrogen Receptor Antagonists |
| Estrogen Receptor Modulators | Selective Estrogen Receptor Modulators |
| Insulin Antagonists | Leukotriene Antagonists |
| Mineralocorticoid Receptor Antagonists | Prostaglandin Antagonists |
| Puberty Inhibitors | Steroid Synthesis Inhibitors |
| 14-alpha Demethylase Inhibitors | 5-alpha Reductase Inhibitors |
| Aromatase Inhibitors | Hormones |
| Anabolic Agents | Androgens |
| Cannabinoid Receptor Modulators | Cannabinoid Receptor Agonists |
| Cannabinoid Receptor Antagonists | Estrogens |
| Estrogens, Non-Steroidal | Phytoestrogens |
| Glucocorticoids | Incretins |
| Mineralocorticoids | Progestins |

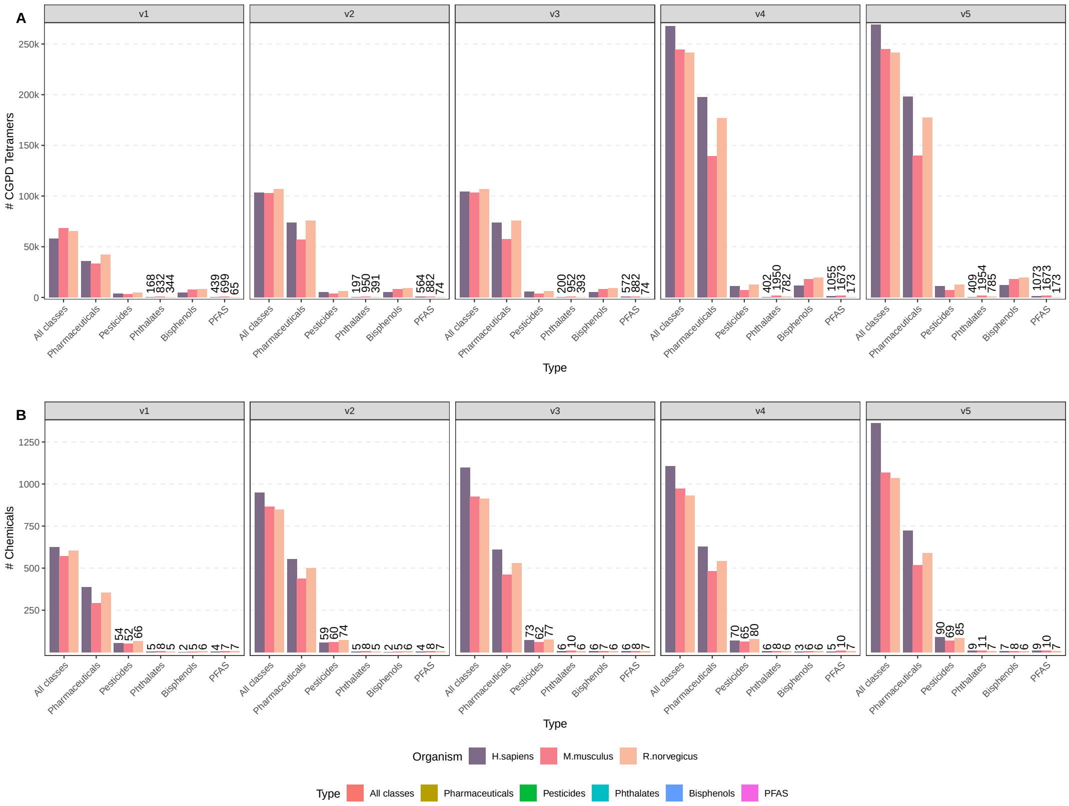


Fig. S2: **Subsetting CTD-derived data to different chemical classes.** (A) Number of CGPD tetramers calculated from `CTDbase`. (B) Number of chemicals for which CGPD tetramers could be calculated. tetramers have been individually calculated for the three target organisms human, rat, and mouse, and five individual filtering strategies - facets 'v1' to 'v5'.. Pesticides have been collected from the EFSA PARAM catalogue (denoted as EFSA Pesticides), from four different CAG reports (EFSA CAGs) and the `CTDbase` itself based on the MeSH dictionary (CTD Pesticides). Pharmaceuticals have been collected from the Drugbank and the `CTDbase` itself based on the MeSH dictionary. Bisphenols, Phthalates, and PFAS have been collected from `CTDbase` using the MeSH dictionary. For details see Section **??**

#### S1 Coverage of tissues in `CTDbase`

*CGPD tetramers per tissue*

*Chemicals per tissue*

##### S1.1 Distribution of CGPD tetramers and Chemicals across tissue groups

The distribution of CGPD tetramers and their underlying unique compounds provides insights into the tissue-specific and organism-specific representation of pesticides, pharmaceuticals, and bisphenols.

For **pesticides**, the nervous system consistently had the highest number of tetramers across all organisms, with rats having the largest count (5027 tetramers) compared to humans (3356) and mice (2824). This pattern was reflected in the number of unique pesticide compounds, with 44 in rats, 32 in humans, and 24 in mice. Additional tissues with substantial pesticide representation included the reproductive system in rats (1914 tetramers, 34 compounds) and the liver across all species.

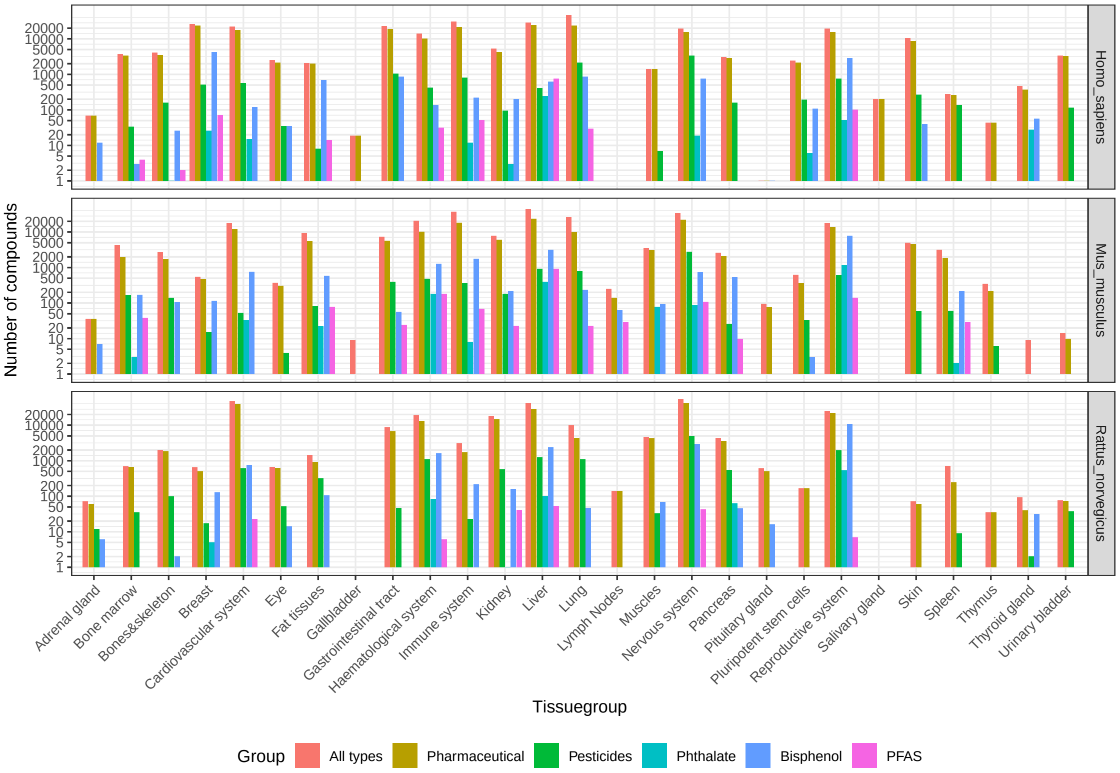


Fig. S3: Impacted tissues by different groups of chemicals and target organism. The number of CGPD tetramers that were calculated for a specific tissuegroup is shown for filtering strategy v5.

**Pharmaceuticals** were extensively represented, with tetramers most abundant in the liver for humans (24179) and mice (23513) and in the nervous system for rats (41931). Unique compounds showed similar patterns, with the nervous system and liver consistently exhibiting high diversity across all organisms. Rats showed the highest chemical diversity in the nervous system (278 compounds), while humans and mice had comparable diversity in the liver and immune system.

**Bisphenols** showed fewer tetramers compared to pesticides and pharmaceuticals, with the reproductive system being the most frequently represented tissue group across all organisms (e.g., 10733 tetramers in rats). Despite the high number of tetramers, the number of unique bisphenol compounds was limited, ranging from 2 to 9 across tissues and organisms. Key tissues included the liver, fat tissues, and cardiovascular system, though their contributions were modest compared to other chemical classes.

These results emphasize significant differences in CGPD tetramer counts and unique compound diversity across chemical classes, tissues, and organisms. While pharmaceuticals demonstrated the highest coverage and diversity, pesticides and bisphenols were more limited in scope, particularly for unique compounds.

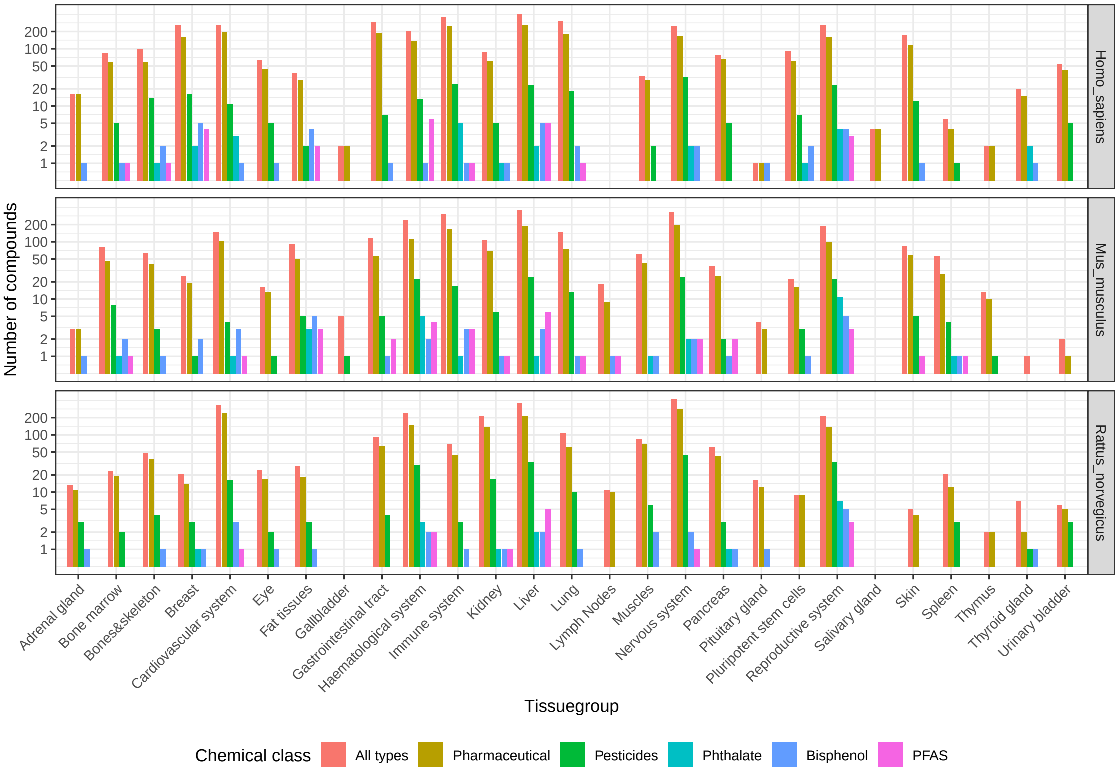


Fig. S4: Impacted tissues by different groups of chemicals and target organism. The number of compounds that effect a specific tissuegroup is shown for filtering strategy v5.

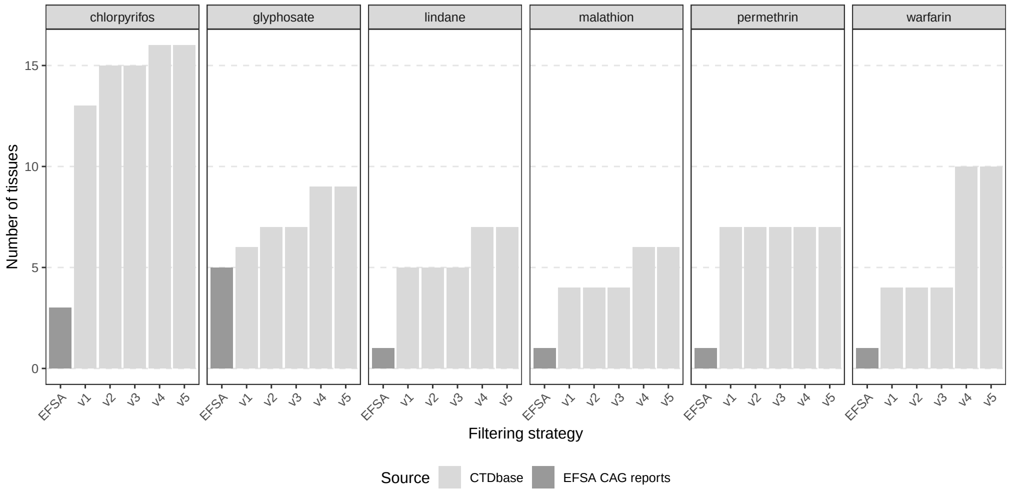


Fig. S5: Comparison of the number of affected tissues for six compounds with high discrepancy between EFSA CAG reports and `CTDbase`. The number of tissues is shown based on the original EFSA CAG report from 2012 and the five different filtering strategies to calculate CGPD tetramers (v1 to v5).

#### S2 Tissue-specific comparisons of EFSA CAG pesticides and `CTD` data

do we need the text? - not referenced in the text yet

The six pesticides with the most significant discrepancies in tissue associations between the EFSA CAG reports and `CTDbase` include 2,4-D, metiram, thiamethoxam, ziram, chlorpyrifos, glyphosate, and warfarin. These pesticides highlight key differences in the number of tissues identified using CGPD tetramers compared to the EFSA CAG definitions.

Pesticides such as 2,4-D, metiram, thiamethoxam, and ziram were linked to a substantial number of tissues in the EFSA CAG reports (10, 7, 8, and 8, respectively) but only one or two tissues using `CTD` data. Conversely, for pesticides such as chlorpyrifos, glyphosate, and warfarin, we identified far more impacted tissues using CGPD tetramers than reported in the EFSA CAGs. Chlorpyrifos, for instance, was associated with 16 tissues in `CTDbase`, compared to just three in the EFSA CAG reports (eye, nervous system, and reproductive system). Glyphosate and warfarin were linked to 9 and 10 tissues, respectively, using CGPD tetramers, compared to 5 and 1 tissue in the EFSA reports.

Supplementary Figure S5 illustrates how different filtering strategies (v1–v5) influenced the identification of tissues for these six pesticides. Notably, warfarin exhibited the most dramatic increase in the number of impacted tissues under filtering strategy v5 compared to v1. This increase was primarily due to the omission of tissue specificity for disease interactions in v5, leading to associations with diseases like coronary artery disease and blood coagulation disorders. These diseases were linked to various tissues, including lung, muscles, spleen, and reproductive organs, based on `CTD`-annotated interactions.

Chlorpyrifos also showed an increase in the number of affected tissues from 13 in v1 to 16 in v5. In contrast, the EFSA CAG reports linked chlorpyrifos to only three tissues. The EFSA CAG classifications relied on data from pesticide databases, Draft Assessment Reports (DARs), and literature published before 2010, identifying approximately 2200 papers for chlorpyrifos. Of these, 169, 33, and 31 papers were linked to the liver, immune system, and kidney, respectively. It remains unclear whether these papers influenced the EFSA CAG focus on three target tissues or were disregarded due to insufficient evidence.

For liver associations, our CGPD tetramer approach linked chlorpyrifos to liver-specific diseases such as liver cirrhosis, fatty liver, hepatomegaly, and chemical-induced liver injury. These associations were supported by manually curated disease interactions with direct evidence, primarily from publications after 2017. An earlier study by Khan (2006) linked chlorpyrifos to ”liver

Table S5: Pesticides that were linked to the nervous system using CTD-derived data. Those compounds are mentioned in the CAG reports but not found to be toxic to the nervous system.

| Chemical | MeSH ID | Chemical | MeSH ID |
|---|---|---|---|
| azoxystrobin | C087670 | carbendazim | C006698 |
| Chlordan | D002706 | cyprodinil | C108338 |
| Diquat | D004178 | fenpyroximate | C415144 |
| glyphosate | C010974 | mepanipyrim | C104083 |
| pyrachlostrobin | C513428 | pyrimethanil | C108337 |
| Warfarin | D014859 | | |

Table S6: Pesticides that were linked to the liver using CTD-derived data. Those compounds are mentioned in the CAG reports but not found to be toxic to the liver.

| Chemical | MeSH ID | Chemical | MeSH ID |
|---|---|---|---|
| 1,2,4-triazole | C045575 | bifenthrin | C099952 |
| Chlorpyrifos | D004390 | cyproconazole | C093628 |
| Diazinon | D003976 | Fenitrothion | D005278 |
| Fenthion | D005284 | fenvalerate | C017690 |
| Hexachlorocyclohexane | D001556 | Monocrotophos | D008999 |
| sedaxane | C583365 | Parathion | D010278 |
| Permethrin | D026023 | vinclozolin | C025643 |
| Warfarin | D014859 | | |

Table S7: Pesticides that were linked to the reproductive system using CTD-derived data. Those compounds are mentioned in the CAG reports but not found to be toxic to the reproductive system.

| Chemical | MeSH ID | Chemical | MeSH ID |
|---|---|---|---|
| bifenthrin | C099952 | Diquat | D004178 |
| Endosulfan | D004726 | Fenitrothion | D005278 |
| Hexachlorocyclohexane | D001556 | Malathion | D008294 |
| Permethrin | D026023 | vinclozolin | C025643 |

Table S8: Pesticides that were linked to the haematological system using CTD-derived data. Those compounds are mentioned in the CAG reports but not found to be toxic to the haematological system.

| Chemical | MeSH ID | Chemical | MeSH ID |
|---|---|---|---|
| acephate | C001969 | bifenthrin | C099952 |
| carbendazim | C006698 | Chlorpyrifos | D004390 |
| cyhalothrin | C037304 | cypermethrin | C017160 |
| decamethrin | C017180 | Diazinon | D003976 |
| Dimethoate | D004117 | Diquat | D004178 |
| Endosulfan | D004726 | enilconazole | C017435 |
| ethion | C100038 | Fenitrothion | D005278 |
| fenvalerate | C017690 | glyphosate | C010974 |
| imidacloprid | C082359 | Malathion | D008294 |
| Permethrin | D026023 | sulfoxaflor | C560328 |
| Thiamethoxam | D000077922 | Warfarin | D014859 |

disease," but the report focused on protective effects of black tea extracts, possibly excluding it from the EFSA CAG report consideration.

Similarly, unspecific diseases such as fibrosis, necrosis, and edema were linked to the kidney based on recent publications, forming CGPD tetramers for chlorpyrifos in this organ. Direct evidence associating chlorpyrifos with "kidney diseases" was supported by four papers published from 2017 onwards.

These findings underscore the potential of CGPD tetramers to provide updated and more comprehensive tissue-specific pesticide associations based on recent curated evidence. They highlight discrepancies in historical datasets and the value of integrating newer data for refining cumulative risk assessments.

Table S9: Phenotypes associated with the chemical cluster linked with Oligospermia. Here, we listed the number of unique chemical-phenotype interactions that are present in this cluster.

| Phenotype name | Phenotype ID | # Chemical-Phenotype Interactions |
|---|---|---|
| spermatogenesis | GO:0007283 | 30 |
| male gonad development | GO:0008584 | 6 |
| ovarian follicle development | GO:0001541 | 6 |
| antral ovarian follicle growth | GO:0001547 | 4 |
| positive regulation of apoptotic process | GO:0043065 | 4 |
| regulation of DNA-templated transcription | GO:0006355 | 3 |
| Leydig cell differentiation | GO:0033327 | 2 |
| positive regulation of epithelial cell proliferation | GO:0050679 | 2 |
| regulation of transcription by RNA polymerase II | GO:0006357 | 2 |
| Sertoli cell proliferation | GO:0060011 | 1 |
| cell population proliferation | GO:0008283 | 1 |
| decidualization | GO:0046697 | 1 |
| negative regulation of cell population proliferation | GO:0008285 | 1 |
| negative regulation of hydrolase activity | GO:0051346 | 1 |
| nitric oxide biosynthetic process | GO:0006809 | 1 |
| positive regulation of autophagy | GO:0010508 | 1 |
| positive regulation of nitric oxide biosynthetic process | GO:0045429 | 1 |
| response to estradiol | GO:0032355 | 1 |
| uterus development | GO:0060065 | 1 |

Table S10: GO enrichment for eight genes associated with the cluster that was linked to the disease hyperglycemia in the haematological system in rats. The clusters were associated with the phenotypes glucose homeostasis (GO:0042593) and the positive regulation of insulin secretion (GO:0032024) and formed in filtering strategies v2 to v5. The eight genes were Gck (glucokinase), Il6 (interleukin 6), Ins1 (Insulin 1), Ins2 (insulin 2), Lepr (leptin receptor), Insr (insulin receptor), Lep (leptin), and Adipoq (adiponectin). We selected only those GO terms with a GeneRatio of 1, meaning that all 8 genes had to be present in the particular GO term.

| GO term | Description | Count | Genes | GeneRatio | GO size | Go ratio |
|---|---|---|---|---|---|---|
| GO:0006006 | glucose metabolic process | 8 | 8 | 1 | 104 | 0.0769 |
| GO:0019318 | hexose metabolic process | 8 | 8 | 1 | 107 | 0.0748 |
| GO:0005996 | monosaccharide metabolic process | 8 | 8 | 1 | 113 | 0.0708 |
| GO:0033500 | carbohydrate homeostasis | 8 | 8 | 1 | 170 | 0.0471 |
| GO:0042593 | glucose homeostasis | 8 | 8 | 1 | 170 | 0.0471 |
| GO:0005975 | carbohydrate metabolic process | 8 | 8 | 1 | 181 | 0.0442 |
| GO:0032870 | cellular response to hormone stimulus | 8 | 8 | 1 | 356 | 0.0225 |
| GO:0051050 | positive regulation of transport | 8 | 8 | 1 | 407 | 0.0197 |
| GO:0048878 | chemical homeostasis | 8 | 8 | 1 | 427 | 0.0187 |
| GO:0044281 | small molecule metabolic process | 8 | 8 | 1 | 474 | 0.0169 |

#### S3 Cluster characterization

Table S11: Chemicals that were grouped and clustered in the rat haematological system. The group was linked with the disease hyperglycemia (MeSH: D006943) and the phenotypes glucose homeostasis (GO:0042593) and positive regulation of insulin secretion (GO:0032024). The 19 chemicals in the top were grouped using the filtering strategies v1 to v5. the other 12 chemicals at the bottom were grouped in this cluster in filtering strategies v2 to v5. A detailed list of the parameter settings is shown in Supplementary Table S12. The column 'Effect' denotes the effect of the compound w.r.t. blood glucose levels. A positive effects leads to decreasing and a negative effect leads to increasing glucose levels.

| Filter | Chemical | MeSH ID | Group | Effect |
|---|---|---|---|---|
| v1 - v5 | 2,2',4,4'-tetrabromodiphenyl ether | C511295 | Flame retardant | negative |
| | Acrylamide | D020106 | Organic chemical | negative |
| | Arsenic | D001151 | Heavy metal | negative |
| | Arsenic Trioxide | D000077237 | Pharmaceutical | negative |
| | Atorvastatin | D000069059 | Pharmaceutical | negative |
| | bisphenol A | C006780 | Bisphenol, Pharmaceutical | negative |
| | Blood Glucose | D001786 | Carbohydrate | negative |
| | Chlorpyrifos | D004390 | Pesticide | negative |
| | dapagliflozin | C529054 | Pharmaceutical | positive |
| | decabromobiphenyl ether | C010902 | Flame retardant | negative |
| | Dexamethasone | D003907 | Pharmaceutical | negative |
| | Dietary Fats | D004041 | Lipid | negative |
| | Fructose | D005632 | Carbohydrate, Pharmaceutical | negative |
| | Glucose | D005947 | Carbohydrate, Pharmaceutical | negative |
| | imidacloprid | C082359 | Pesticide | negative |
| | Letrozole | D000077289 | Pharmaceutical | negative |
| | Nicotine | D009538 | Pesticide | negative |
| | sodium arsenite | C017947 | Pharmaceutical | negative |
| | Streptozocin | D013311 | Pharmaceutical | negative |
| v2 - v5 | acephate | C001969 | Pesticide | negative |
| | cobaltiprotoporphyrin | C007095 | Heterocyclic compounds | positive |
| | corilagin | C049096 | Organic chemical | positive |
| | Curcumin | D003474 | Pharmaceutical | positive |
| | Diazinon | D003976 | Pesticide | negative |
| | Fatty Acids, Omega-3 | D015525 | Lipid, Pharmaceutical | negative |
| | Glyburide | D005905 | Pharmaceutical | positive |
| | Lead | D007854 | Heavy metal | negative |
| | Metformin | D008687 | Pharmaceutical | positive |
| | Propolis | D011429 | Mixture, Antimicrobial | positive |
| | Quercetin | D011794 | Pharmaceutical | positive |
| | Resveratrol | D000077185 | Pharmaceutical | positive |

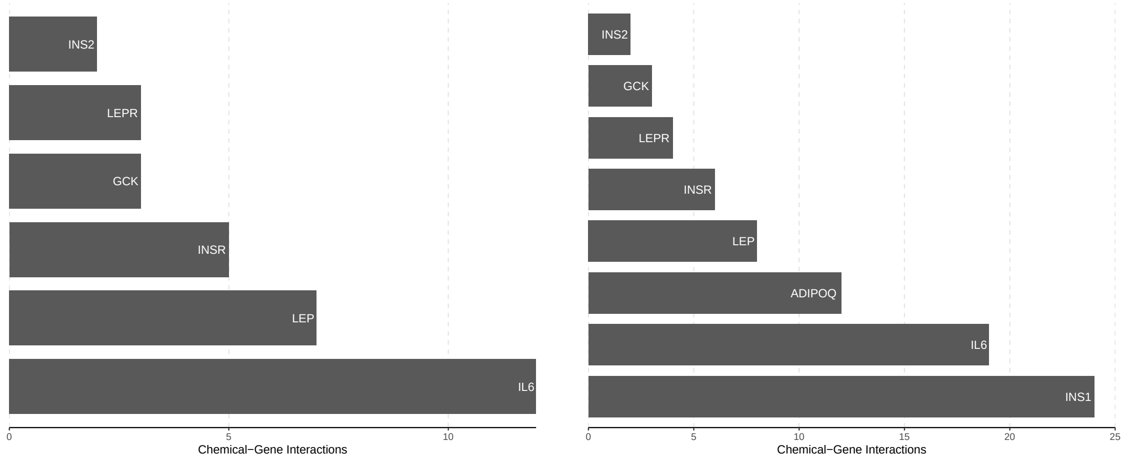


(a) Filtering strategy v1

(b) Filtering strategies v2 - v5

Fig. S6: **Chemical-gene interactions in Hyperglycemia associated cluster.** (a) Unique chemical-gene interactions for the six associated genes in the chemical cluster formed in filtering strategy v1 containing 19 chemicals. (a) Unique chemical-gene interactions for the eight associated genes in the chemical cluster formed in filtering strategies v2-v5 containing 31 chemicals.

Table S12: Clusters containing chemicals that were linked to the disease hyperglycemia (MeSH: D006943) in the Haematological system of *Rattus norvegicus*. In total, 40 different parameter settings in filtering strategies v2 to v5 resulted in clusters with the same set of 31 chemicals. 10 different parameter settings in filtering strategy v1 resulted in a reduced cluster with 19 chemicals. The chemicals are listed in Supplementary Table S11

| Filter | Grouping | Phenotype clustering | | Chemical clustering | | Chemicals |
|---|---|---|---|---|---|---|
| | | Applied | Cutoff | Applied | Cutoff | |
| v1 | pd | - | - | x | 0.75 | 19 |
| v1 | pd | - | - | x | 0.9 | 19 |
| v1 | pd | x | 0.75 | x | 0.5 | 19 |
| v1 | pd | x | 0.75 | x | 0.75 | 19 |
| v1 | pd | x | 0.75 | x | 0.9 | 19 |
| v1 | pd | x | 0.9 | x | 0.5 | 19 |
| v1 | pd | x | 0.9 | x | 0.75 | 19 |
| v1 | pd | x | 0.9 | x | 0.9 | 19 |
| v1 | pd | x | 0.75 | - | - | 19 |
| v1 | pd | x | 0.9 | - | - | 19 |
| v2 | pd | - | - | x | 0.75 | 31 |
| v2 | pd | - | - | x | 0.9 | 31 |
| v2 | pd | x | 0.75 | x | 0.5 | 31 |
| v2 | pd | x | 0.75 | x | 0.75 | 31 |
| v2 | pd | x | 0.75 | x | 0.9 | 31 |
| v2 | pd | x | 0.9 | x | 0.5 | 31 |
| v2 | pd | x | 0.9 | x | 0.75 | 31 |
| v2 | pd | x | 0.9 | x | 0.9 | 31 |
| v2 | pd | x | 0.75 | - | - | 31 |
| v2 | pd | x | 0.9 | - | - | 31 |
| v3 | pd | - | - | x | 0.75 | 31 |
| v3 | pd | - | - | x | 0.9 | 31 |
| v3 | pd | x | 0.75 | x | 0.5 | 31 |
| v3 | pd | x | 0.75 | x | 0.75 | 31 |
| v3 | pd | x | 0.75 | x | 0.9 | 31 |
| v3 | pd | x | 0.9 | x | 0.5 | 31 |
| v3 | pd | x | 0.9 | x | 0.75 | 31 |
| v3 | pd | x | 0.9 | x | 0.9 | 31 |
| v3 | pd | x | 0.75 | - | - | 31 |
| v3 | pd | x | 0.9 | - | - | 31 |
| v4 | pd | - | - | x | 0.75 | 31 |
| v4 | pd | - | - | x | 0.9 | 31 |
| v4 | pd | x | 0.75 | x | 0.5 | 31 |
| v4 | pd | x | 0.75 | x | 0.75 | 31 |
| v4 | pd | x | 0.75 | x | 0.9 | 31 |
| v4 | pd | x | 0.9 | x | 0.5 | 31 |
| v4 | pd | x | 0.9 | x | 0.75 | 31 |
| v4 | pd | x | 0.9 | x | 0.9 | 31 |
| v4 | pd | x | 0.75 | - | - | 31 |
| v4 | pd | x | 0.9 | - | - | 31 |
| v5 | pd | - | - | x | 0.75 | 31 |
| v5 | pd | - | - | x | 0.9 | 31 |
| v5 | pd | x | 0.75 | x | 0.5 | 31 |
| v5 | pd | x | 0.75 | x | 0.75 | 31 |
| v5 | pd | x | 0.75 | x | 0.9 | 31 |
| v5 | pd | x | 0.9 | x | 0.5 | 31 |
| v5 | pd | x | 0.9 | x | 0.75 | 31 |
| v5 | pd | x | 0.9 | x | 0.9 | 31 |
| v5 | pd | x | 0.75 | - | - | 31 |
| v5 | pd | x | 0.9 | - | - | 31 |
